# Developmental disruption of *Mef2c* in Medial Ganglionic Eminence-derived cortical inhibitory interneurons impairs cellular and circuit function

**DOI:** 10.1101/2024.05.01.592084

**Authors:** Claire Ward, Kaoutsar Nasrallah, Duy Tran, Ehsan Sabri, Arenski Vazquez, Lucas Sjulson, Pablo E. Castillo, Renata Batista-Brito

**Affiliations:** Dominick P. Purpura Department of Neuroscience, Albert Einstein College of Medicine, Bronx, NY 10461; Department of Biological Sciences, Fordham University, Bronx, NY 10458; Department of Psychiatry and Behavioral Sciences, Albert Einstein College of Medicine, Bronx, NY 10461; Department of Genetics, Albert Einstein College of Medicine, Bronx, NY 10461

## Abstract

*MEF2C* is strongly linked to various neurodevelopmental disorders (NDDs) including autism, intellectual disability, schizophrenia, and attention-deficit/hyperactivity. Mice constitutively lacking one copy of *Mef2c*, or selectively lacking both copies of *Mef2c* in cortical excitatory neurons, display a variety of behavioral phenotypes associated with NDDs. The MEF2C protein is a transcription factor necessary for cellular development and synaptic modulation of excitatory neurons. MEF2C is also expressed in a subset of cortical GABAergic inhibitory neurons, but its function in those cell types remains largely unknown. Using conditional deletions of the *Mef2c* gene in mice, we investigated the role of MEF2C in Parvalbumin-expressing Interneurons (PV-INs), the largest subpopulation of cortical GABAergic cells, at two developmental timepoints. We performed slice electrophysiology, *in vivo* recordings, and behavior assays to test how embryonic and late postnatal loss of MEF2C from GABAergic interneurons impacts their survival and maturation, and alters brain function and behavior. We found that loss of MEF2C from PV-INs during embryonic, but not late postnatal, development resulted in reduced PV-IN number and failure of PV-INs to molecularly and synaptically mature. In association with these deficits, early loss of MEF2C in GABAergic interneurons lead to abnormal cortical network activity, hyperactive and stereotypic behavior, and impaired cognitive and social behavior. Our findings indicate that MEF2C expression is critical for the development of cortical GABAergic interneurons, particularly PV-INs. Embryonic loss of function of MEF2C mediates dysfunction of GABAergic interneurons, leading to altered *in vivo* patterns of cortical activity and behavioral phenotypes associated with neurodevelopmental disorders.

## Introduction

MEF2C is a transcription factor widely associated with various neurodevelopmental disorders (NDDs) including autism, intellectual disability, schizophrenia, and attention-deficit/hyperactivity disorder (1-6). MEF2C is broadly expressed in cortical excitatory neurons, where it regulates synaptic transmission and is involved in experience-dependent synaptic remodeling (2, 3, 5). MEF2C is also expressed in cortical GABAergic interneurons, particularly parvalbumin-expressing cortical interneurons (PV-INs). Heterozygous MEF2C loss-of-function mutations in humans result in MEF2C haploinsufficiency syndrome, which is characterized by various autism spectrum disorder (ASD) symptoms, including poor eye contact, absence of speech, intellectual disability, seizures, stereotypic movements, and hyperactivity (2, 4, 6).

MEF2C haploinsufficiency mouse models show that developmental loss of one copy of *Mef2c* across all cell types leads to ASD-associated behaviors (1, 7). At a cortical level, these mice have increased excitatory to inhibitory (E/I) neurotransmission, suggesting reduced inhibition (7). In contrast, conditional removal of two copies of *Mef2c* in Emx1-expressing cortical excitatory neurons results in reduced E/I neurotransmission and leads to some, but not all, of the behavioral phenotypes observed in the MEF2C haploinsufficiency mouse model (3). Taken together, these studies suggest that loss of *Mef2c* in cortical GABAergic INs might underlie some of the circuit and behavioral phenotypes observed in the MEF2C haploinsufficiency model.

## Results

### Deletion of *Mef2c* from interneuron progenitors leads to both PV-IN cell loss and impaired PV-IN maturation

Previous studies have shown that embryonic *Mef2c* expression is enriched in the PV-IN lineage, a subset of INs that originates in the medial ganglionic eminence (MGE), and is the earliest predictor of PV-IN fate (9). We have fate mapped MGE-derived INs, PV-INs and somatostatin expressing Interneurons (SST-INs), by crossing the PV-Cre and SST-Cre driver mouse lines to a mouse line with Cre-dependent expression of a fluorescent reporter (RCE, GFP expression). INs derived from the caudal ganglionic eminence (CGE) where labeled in the same manner using a 5HT3a-Cre driver line. We show that MEF2C is expressed in virtually all neocortical fate-mapped PV-INs (98.0 ± 0.7%) at postnatal day 25 (P25) cortex, and in a subset of SST-INs (24.4 ± 2.3%) (Figure 1A-B) (9, 10). In contrast, *Mef2c* is largely excluded from CGE-derived INs (Figure 1A-B).

**Figure 1.**
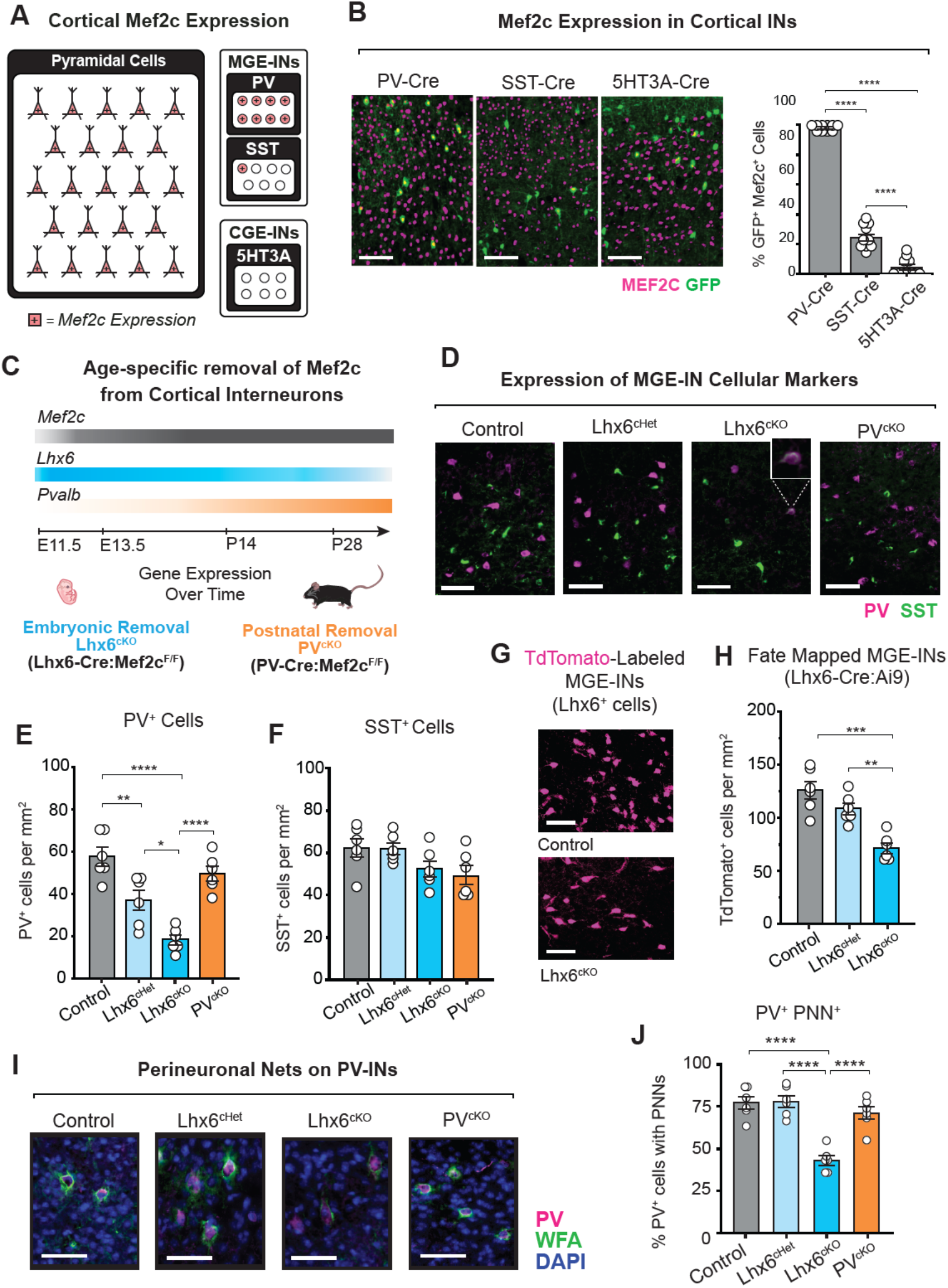
Removal of *Mef2c* from MGE-progenitors leads to a reduction in PV^+^ neurons. **(A)** Schematic depicting *Mef2c* expression among major classes of cortical neurons (red), including excitatory pyramidal cells, inhibitory interneurons (INs) derived from the MGE (PV^+^, SST^+^), and CGE (5HT3A^+^). **(B)** Representative images and quantification of *Mef2c* expression in IN populations in the primary visual cortex (V1) of postnatal day 25 (P25) animals. Cells were genetically labeled through crossing transgenic Cre and (PV-Cre, SST-Cre, 5HT3A-Cre) Cre-reporter (RCE, GFP expression) mouse lines. Colocalization (yellow) of GFP^+^ cells with immunostaining for MEF2C (magenta). White scalebars = 300 μm. For all groups n=10 (3 animals, 2-5 slices per animal). **(C)** Transgenic mouse lines used to assess the early and late roles of *Mef2c* in PV-IN development and function based on the expression profiles of *Mef2c, Lhx6*, and *Pvalb* (Allen Brain Institute Developing Mouse Brain Atlas ISH). **(D)** Representative images of PV (magenta) and SST (green) ex-pression in animals with embryonic removal of either one or both *Mef2c* alleles (Lhx6^cHet^, Lhx6^cKO^) or postnatal removal of *Mef2c* (PV^cKO^) compared to control animals. Inset in Lhx6^cKO^ panel shows an example cell positive for both SST and PV (yellow). White scalebars = 200 μm. **(E, F)** Quantification of PV^+^ cells and SST^+^ cells in V1 of P25 animals normalized by area (cells/mm^2^). For all groups n=6 (3 animals, 1-3 sections per animal). **(G, H)** Representative images of MGE-INs (magenta) in V1 of P25 animals. Cells were genetically labeled through crossing Lhx6-Cre and Cre-reporter mouse lines with wild-type or floxed *Mef2c* alleles (Ai9, TdTomato expression). White scalebars = 200μm. **(H)** Number of TdTomato-expressing cells normalized by area (cells/mm^2^). For all groups n=6 (3 animals, 1-3 sections per animal). **(I, J)** Staining and quantification of perineuronal nets (PNN, green) around PV^+^ cells (magenta) in V1 of P25 animals. White scalebar = 200μm. **(J)** Percent of PV^+^ cells with PNN stain-ing normalized to the total number of PV^+^ cells. For all groups n=6 (3 animals, 1-3 sections per animal). Across all plots: One-way ANOVA with Tukey HSD post-test. *p<0.05, **p<0.01, ***p<0.001, **** p<0.0001. Each point corresponds to values from a single 10x field of view. Data are pre-sented as mean ± SEM. Detailed statistics in Table S1.

Considering that *Mef2c* is expressed in both developing and mature cortical IN populations, we used transgenic mouse lines to conditionally remove *Mef2c* from MGE-INs in a time-specific manner. We generated Lhx6-Cre:*Mef2c*^F/+^ (Lhx6^cHet^) and Lhx6-Cre:*Mef2c*^F/F^ (Lhx6^cKO^) mice in order to remove one or two copies of *Mef2c* from MGE-INs during embryonic development (11-16). We generated PV-Cre:*Mef2c*^F/F^ (PV^cKO^) mice in order to remove two copies of *Mef2c* from PV-INs past P15 (17-20). We assessed whether disruption of *Mef2c* in embryonic development would lead to impaired interneuron survival and maturation through comparing the number of PV^+^ and SST^+^ cells in both primary sensory and associative cortical regions (Figure 1C, D). We found that embryonic removal of *Mef2c* diminishes the number of cells immunolabeled with PV in both V1 and ACC, with an approximate two-thirds reduction in Lhx6^cKO^ animals compared to control animals (Figure 1E, Figure S1B). Within V1, embryonic loss of only one *Mef2c* allele (Lhx6^cHet^) resulted in an intermediate reduction of PV^+^ cells at P25. In contrast to early removal, postnatal loss of *Mef2c* in PV-INs in PV^cKO^ mice did not significantly reduce the numbers of PV-expressing cells (Figure 1E, Figure S1B). In all groups, the number of SST-expressing cells did not change in either V1 or ACC (Figure 1F, Figure S1C). We also observed decrease in PV^+^ cells within the dentate gyrus of the dorsal hippocampus of Lhx6^cKO^ mice (Figure S1I). While populations of PV^+^ and SST^+^ cells do not typically overlap in mice (10, 21-23), a small portion of cells co-expressed SST and PV in Lhx6^cKO^ tissue, supporting the notion that embryonic loss of *Mef2c* may result in a small number of INs that have a mixed fate (Figure 1D inset, Figure S1A inset, Figure S1D-E)(24).

To understand whether the reduction of PV^+^ cells in Lhx6^cKO^ animals was primarily due to reduced number of MGE-INs or simply a downregulation of PV expression, we bred our animals to a Cre-reporter mouse line (Ai9, TdTomato expression). We saw a significant reduction in the number of cortical MGE-INs in P25 Lhx6^cKO^ animals in both V1 and ACC (Figure 1G-H, Figure S1F). We did not see an increase in the number of MGE-INs negative for both PV and SST within Lhx6^cKO^ animals relative to control animals, supporting the notion that the reduction of PV^+^ cells is primarily driven by cell loss (Figure S1F-H). We observed however that PV expression levels might be reduced in Lhx6^cKO^ animals, an effect that could be due to reduced activity of PV-INs (25). Lhx6^cHet^ animals, while not having MGE-IN cell loss (Figure 1H, Figure S1F-H), had reduced number of PV^+^ cells in V1 (Figure 1E). We also assessed the presence of perineuronal nets (PNNs) surrounding PV-INs in V1, as the appearance of these extracellular matrix structures coincides with an increase in PV expression (26-29). We found that embryonic removal of *Mef2c* from MGE-INs leads to a reduction in the presence of PNNs (Figure 1I, J). Because the onset of PV expression is gradual, we confirmed the efficiency of *Mef2c* removal in 4-month-old PV^cKO^ animals, where PV-expression should peak (Figure S1J). We did not see a reduction in the number of PV^+^ cells or a reduction in the percentage of PV^+^ cells with PNNs in adult PV^cKO^ animals (Figure S1K). Taken together, our data shows that embryonic, but not postnatal, removal of *Mef2c* leads to a reduction in the number of neocortical PV-lineage cells.

### Early, but not late, loss of *Mef2c* reduces excitatory drive onto PV-INs

We then investigated whether early and/or late loss of *Mef2c* impairs the synaptic maturation of the remaining PV-INs. Given that MEF2C is known to regulate synapses in cortical excitatory neurons (1-3, 5, 7, 30-32), we tested whether early vs. late loss of *Mef2c* impacts glutamatergic excitatory drive onto PV-INs at adult ages. To this end, we used electrophysiology in acute brain slices obtained from Control, Lhx6^cKO^, Lhx6^cHet^, and PV^cKO^ adult mice (Figure 2). To identify PV-INs, we expressed YFP specifically in these neurons by injecting AAV-S5E2-C1V1-EYFP, a PV-lineage specific virus that targets PV-IN lineage independently of PV expression, into the mouse V1 3-5 weeks prior to electrophysiology recordings (Figure 2A-B) (8). Spontaneous excitatory postsynaptic currents (sEPSCs) were then monitored in labeled PV-INs using whole-cell voltage-clamp recordings (Vholding = -60 mV) in the presence of GABAA and GABAB receptor antagonists (see Methods). Early deletion of *Mef2c* in Lhx6^cKO^ mice led to ∼80% reduction in sEPSC frequency in PV-INs, whereas it did not significantly change sEPSC amplitude (Figure 2C-D). In addition, Lhx6^cHet^ mice showed an intermediate phenotype as compared to control and Lhx6^cKO^ mice, with ∼40% reduction in sEPSCs frequency and no significant change in amplitude (Fig 2C-D). In contrast, late loss of *Mef2c* in PV^cKO^ did not impact sEPSC frequency or amplitude into PV-INs (Figure C-D). These results strongly suggest that early, but not late, *Mef2c* expression is necessary for developing normal excitatory drive onto PV-INs.

**Figure 2.**
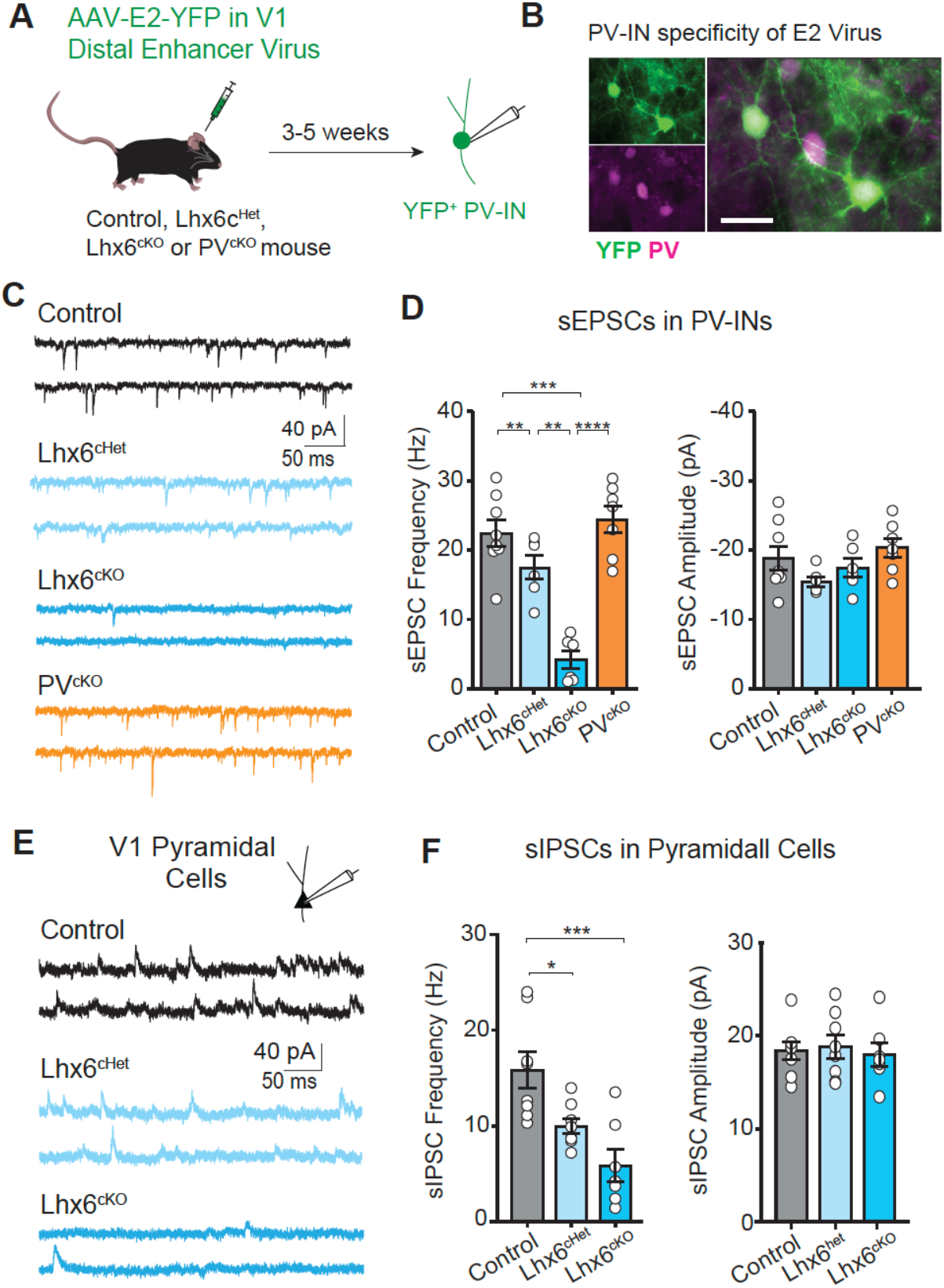
Embryonic removal of *Mef2c* from MGE-INs impairs excitatory transmission onto PV-INs. **(A-D)** Whole cell recordings obtained from YFP-expressing PV-INs (Vh = -60 mV) in V1 cortex of control (*Mef2c*^+^/^+^), Lhx6^cKO^, Lhx6^cHet^ and PV^cKO^ mice. **(A-B)** AAV-PV(En)-YFP was injected into primary visual cortex (V1) to express YFP specifically in PV-INs (white scalebar = 50μm). **(C-D)** Single traces and summary plots of spontaneous EPSC (sEPSC) frequency and amplitude in controls animals (n = 8, 4 mice) compared Lhx6^cHet^ (n = 6, 3 mice), Lhx6^cKO^ (n = 6, 3 mice) and PV^cKO^ (n = 7, 2 mice). **(E-F)** Spontaneous IPSCs (sIPSCs) recordings from pyramidal neurons in V1 of control, Lhx6^cHet^, and Lhx6^cKO^ animals. Representative traces **(E)** and summary plots **(F)** of sIPSC frequency and amplitude in controls animals (n = 8, 5 mice) compared Lhx6^cHet^ (n = 8, 4 mice), Lhx6^cKO^ (n = 7, 3 mice). Across all plots: One-way ANOVA with Tukey HSD post-test. *p<0.05, **p<0.01, ***p<0.001, **** p<0.0001. Each point corresponds to a single patched cell. Data are presented as mean ± SEM. Detailed statistics in Table S3. Across all plots: One-way ANOVA with Tukey HSD post-test. *p<0.05, **p<0.01, ***p<0.001, **** p<0.0001. Each point corresponds to a single patched cell. Data are presented as mean ± SEM. Detailed statistics in Table S3.

### Early *Mef2c* loss in MGE-derived GABAergic neurons reduces inhibitory synaptic transmission onto principal cells

Given the reduced number of PV-INs (Figure 1E) in Lhx6^cKO^ and the fact PV-INs have synaptic deficits in Lhx6^cKO^ and Lhx6^cHet^ mice (Figure 2D), we then tested if GABAergic transmission onto cortical pyramidal cells (PYR) is reduced in adult mutant animals. We recorded spontaneous inhibitory postsynaptic currents (sIPSCs, Vholding = ^+^10 mV) in the presence of AMPA and NMDA receptor antagonists (10 μM NBQX and 50 μM D-APV, respectively). We found that early *Mef2c* deletion in Lhx6^cKO^ mice was accompanied by a robust decrease in sIPSC frequency in PYR but not sIPSC amplitude (Figure 2E-F). In addition, deleting one copy of *Mef2c* in early developmental stages was sufficient to reduce sIPSC frequency in Lhx6^cHet^ mice (Figure 2E-F).

### *Mef2c* loss in MGE-derived GABAergic neurons alters *in vivo* patterns of cortical activity

To examine the effects of early and late *Mef2c* deletion on *in vivo* cortical activity, we performed extracellular recordings of local field potential (LFP), a measure of local network activity, in the V1 and ACC in awake, head-fixed mice (Figure 3, Figure S2). To ensure that comparisons between animals were carried out within similar behavioral states, we restricted the analysis to periods when the animal is active as quantified by locomotion (33). Early *Mef2c* removal in Lhx6^cKO^ mice caused alterations in local network activity both in V1 (Figure 3C-E), and ACC (Figure S2A-C). Lhx6^cKO^ animals showed a decrease in 1-4 Hz LFP power compared to controls, and a robust increase in beta- (12-35 Hz), low gamma- (35-60 Hz), and high gamma-range (60-100 Hz) LFP power in both V1 and ACC of Lhx6^cKO^ animals. Recent analysis focusing on the aperiodic (or non-oscillatory) parameters of LFP signals, has revealed that the power spectrum slope can dynamically change with the levels of inhibition, with an increase in GABAergic tone being reflected in an increase in slope (34, 35). We show that Lhx6^cKO^ animals have significantly reduced slope in comparison with littermate controls in both V1 and ACC (Figure 3E, Figure S2C), further suggestive of reduced inhibition, and in line with the imbalance in excitatory to inhibitory activity that we see at the cellular level (Figure 2). In vivo LFP network activity was not altered in Lhx6^cHet^ or PV^cKO^, indicating that the loss of *Mef2c* leads to abnormal cortical activity in a dose- and age-specific manner (Figure 3F-H, Figure S2D-E).

**Figure 3.**
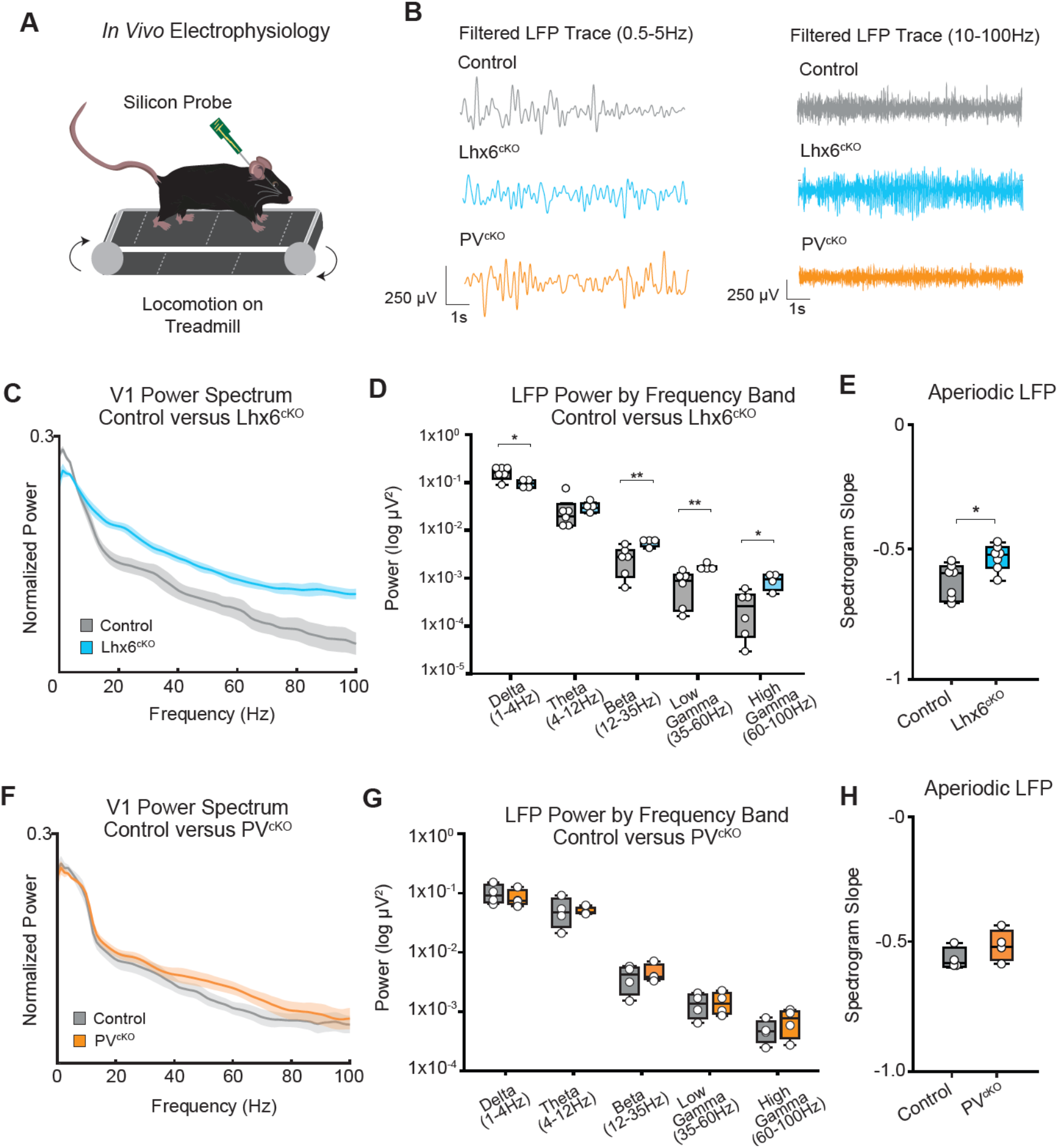
Embryonic loss of *Mef2c* in MGE-INs leads to changes in cortical local field potential activity. **(A)** *In vivo* electrophysiology recordings from primary visual cortex (V1) performed in awake Lhx6^cKO^ mutants and littermate controls. **(B)** Example bandpass-filtered Local Field Potential (LFP) traces within V1 of control (gray), Lhx6^cKO^ (blue), and PV^cKO^ (orange) animals. **(C)** Population-averaged normalized V1 power spectra during locomotion periods for control (6 sessions) and Lhx6^cKO^ animals (4 sessions). **(D)** V1 LFP power compared between control (7 sessions) and Lhx6^cKO^ animals (7 sessions) across frequencies: 1-4 Hz, 4-12 Hz, 12-35 Hz, 35-60 Hz, and 60-100 Hz bands during periods of locomotion. **(E)** Slope of the aperiodic component of the LFP power spectrum. **(F)** Population-averaged normalized V1 power spectra during locomotion periods for control (6 sessions) and PV^cKO^ animals (4 sessions). **(G)** V1 LFP power compared between PV^cKO^ mutants (4 sessions) and control animals (4 sessions) across frequencies: 1-4 Hz, 4-12 Hz, 12-35 Hz, 35-60 Hz, and 60-100 Hz bands during periods of locomotion. **(H)** Slope of the aperiodic component of the LFP power spectrum for PV^cKO^ mutants (4 sessions) and control animals (4 sessions). Student’s t-test: * p < 0.05, ** p < 0.01, *** p< 0.001, ****p<0.0001. Each point corresponds to a single recording session. Data are presented as mean ± SEM. Detailed statistics in Table S4.

### Early, but not late, removal of *Mef2c* leads to hyper locomotor activity and lack of habituation

Given the cellular and network effects of embryonic loss of *Mef2c* from MGE-INs, we examined whether these changes gave rise to NDD-associated behavioral phenotypes. Lhx6^cKO^ mice, but not Lhx6^cHet^ and PV^cKO^ mice, showed increased locomotion in the open field arena (Figure 4A-B). Lhx6^cKO^ animals also exhibited increased jumping behavior (Figure 4C). Unlike all other groups, Lhx6^cKO^ animals did not show reduced locomotion and increased grooming during the latter portion of the assay, suggesting a lack of habituation to a new environment (Figure 4D-E).

**Figure 4.**
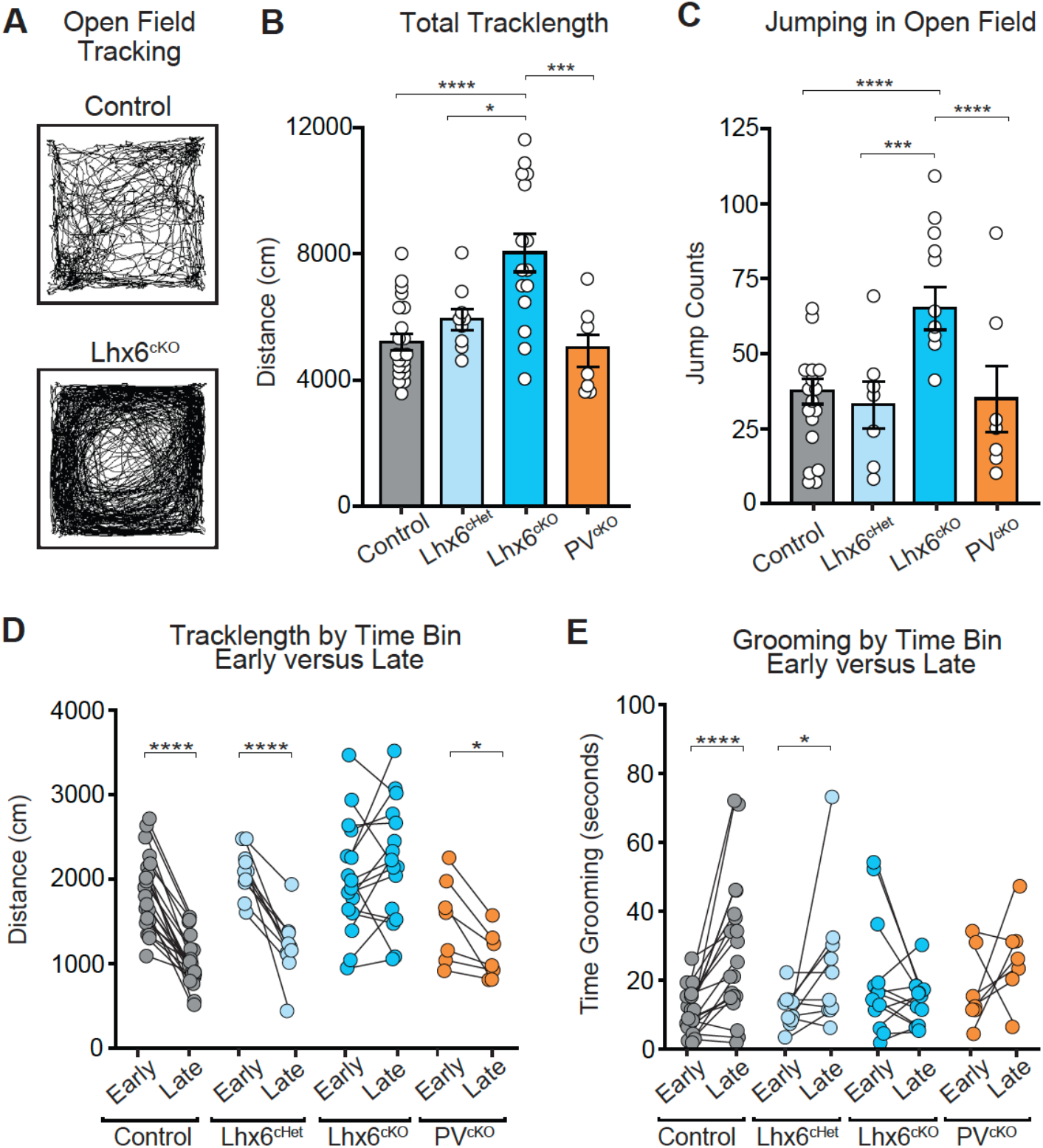
Hyperactivity in mice with embryonic loss of *Mef2c* in MGE-INs. **(A-B)** Representative tracking paths for Control and Lhx6^cKO^ animals in an open field arena. Within a period of 12 minutes, Lhx6^cKO^ (n=15) animals traveled a greater distance (cm) than control (n=21) and Lhx6^cHet^ (n=9), and PV^cKO^ (n=7) animals. **(C)** Increased jumping behavior is seen in Lhx6^cKO^ mutant animals (Control n=17, Lhx6^cHet^ n=7, Lhx6^cKO^ n=10, PV^cKO^ n=7). **(D)** Tracklength binned by the first and last three minutes of the open field assay (early vs. late). During the first three minutes there was no difference in tracklength between animal groups (Control n=21, Lhx6^cHet^ n=10, Lhx6^cKO^ n=15, PV^cKO^ n=6). **(E)** Grooming time in the open field arena compared between early and late bins of the session, where Lhx6^cKO^ mutant animals did not increase grooming behavior toward the end of the session (Control n=20, Lhx6^cHet^ n=10, Lhx6^cKO^ n=13, PV^cKO^ n=7). For plots B and C: One-way ANOVA with Tukey HSD post-test. For plots D and E: Two-way repeated measures ANOVA with Bonferroni post hoc comparison. Each point corresponds to an individual animal. * p < 0.05, ** p < 0.01, *** p< 0.001, ****p<0.0001. Data are presented as mean ± SEM. Detailed statistics in Table S5.

### Early removal of *Mef2c* causes NDD-associated stereotypical movement and increased exploration of elevated plus- and y-maze environments

Paw clasping during tail suspension is a reflex that is most notably present in mouse models of Rett Syndrome (MECP2) but is also observed in other ASD-related mouse models, including animals with disrupted expression of Shank2, Kdm5, and Hdac3, and in the haploinsufficiency model of *Mef2c* (7, 30, 36-43). Lhx6^cKO^ animals exhibited paw clasping, which was not observed in control and Lhx6^cHet^ animals (Figure 5A). Several mouse models of NDDs also have altered exploration of open/closed arms in an elevated plus maze (EPM), an assay that probes the innate conflict between exploration (open arm) and safety (closed arm) (44). Increased open arm exploration is present in previous *Mef2c* animal models (1, 36), and has also been directly linked to altered activity of cortical interneurons (23). We found that Lhx6^cKO^ animals spent significantly more time in the open arms of the maze, while Lhx6^cHet^ animals resembled controls animals (Figure 5C). Similarly, Lhx6^cKO^ animals made more entries into the open arms and had a higher track length than controls, demonstrating that the increased duration in the open arms is likely attributable to more movement (i.e. active exploration) within the open arms rather than the animals simply sitting within the open arms (Figure 5D-E). We next performed a y-maze assay to evaluate whether Lhx6^cKO^ animals exhibited a spontaneous alternation behavior that is innate to wild type animals (Figure 5F). We found that Lhx6^cKO^ animals showed no difference in the percentage of entries that resulted in a complete sequence of entries into each arm of the maze (Figure 5G), however we observed an increase in track length, with little time in the maze center, and more arm entries (Figure 5H-J). Interestingly, Lhx6^cKO^ animals often made continuous movements between arms and were less likely to make partial arm entries than control mice (Figure 5K).

**Figure 5.**
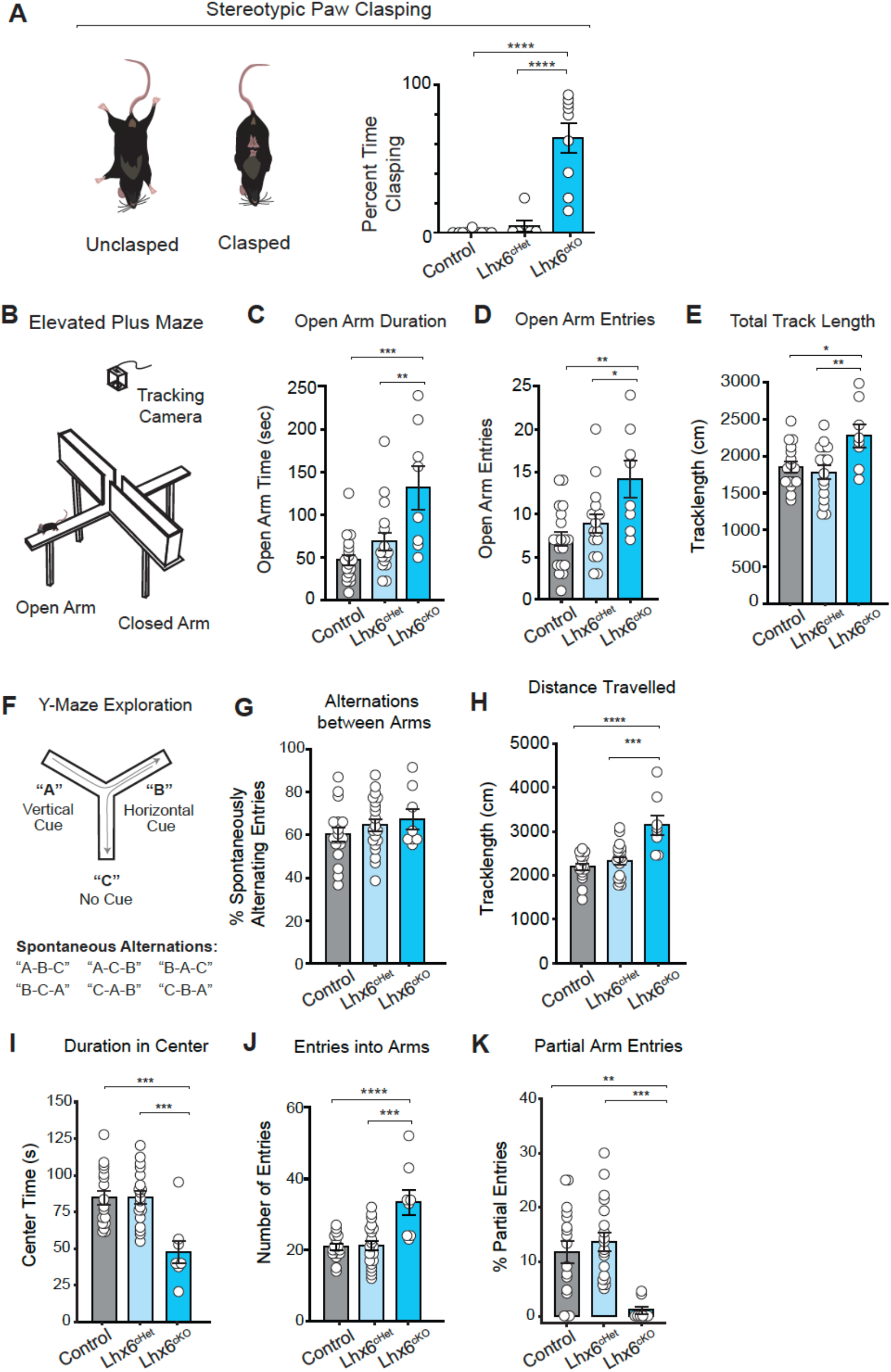
Embryonic loss of *Mef2c* from MGE-progenitors leads to stereotyped paw clasping and atypical exploration in elevated plus maze and y-maze assays. **(A)** Animals were assessed for a paw clasping reflex during a two-minute period of tail suspension. Lhx6^cKO^ animals exhibited high levels of paw clasping, which was not observed in other groups (Control n=13, Lhx6^cHet^ n=8, Lhx6^cKO^ n=11). **(B-E)** Behavior was evaluated in an Elevated Plus Maze assay, in which the animals were tracked to measure the duration and number of entries into the arms of the maze that are open across a 5-minute testing period (Control n=19, Lhx6^cHet^ n=16, Lhx6^cKO^ n=8). **(C-D)** The duration of time spent in the open arms (seconds) and number of open arm entries was higher in Lhx6^cKO^ animals compared to control and Lhx6^cHet^ animals. **(E)** Additionally, the overall tracklength was higher in Lhx6^cKO^ animals compared to control and Lhx6^cHet^ animals. **(F-K)** Behavior was evaluated in a Y-Maze assay, where the sequence of entries into unique maze arms was recorded to determine the number of spontaneous alternations between all three arms (Control n=19, Lhx6^cHet^ n=16, Lhx6^cKO^ n=8). Spontaneous alternations were counted as the number of entries resulting in sequences that contain “A-B-C”, “A-C-B”, “B-A-C”, “B-C-A”, “C-A-B”, and “C-B-A” **(G)** Number of spontaneous alternations normalized by the total number of entries. Lhx6^cKO^ animals carried out the same percentage of spontaneous alternations compared to other groups. **(H-I)** Lhx6^cKO^ animals had higher track lengths and spent less time in the center of the maze. **(J)** Lhx6^cKO^ animals exhibited a higher number of arm entries. **(K)** Lhx6^cKO^ animals performed few partial arm entries, which was counted as a full-body entry into the arm where the animal returned to the maze center before continuing toward the end of one arm. One-way ANOVA with Tukey HSD post-test: * p < 0.05, ** p < 0.01, *** p< 0.001, ****p<0.0001. Each point corresponds to an individual animal. Data are presented as mean ± SEM. Detailed statistics in Table S6.

### Early removal of *Mef2c* causes abnormal social preferences

Deficits in social interaction are a core feature of ASD, and behavioral assays designed to probe the highly social nature of mice have revealed phenotypes common to mouse models of ASD (44, 45). We examined sociability in our animals using the 3-chamber social preference assay, in which animals explored chambers containing social (non-familiar mouse) or non-social stimuli (small object) (Figure 6A). As expected, control and Lhx6^cHet^ animals exhibited greater exploration of the animal stimulus compared to the object stimulus, while Lhx6^cKO^ animals showed no significant difference between the animal and object stimuli (Figure 6B). Because the 3-chamber social preference assay does not allow for direct and reciprocal interactions, we also carried out a social interaction assay within an open field arena where experimental animals were exposed to a novel, conspecific stimulus animal (Figure S3). We observed fewer bouts of social investigation in Lhx6^cKO^ animals compared to both control and Lhx6^cHet^ animals (Figure S3B), which were also shorter in duration (Figure S3C). As expected, based on the open field assay, Lhx6^cKO^ animals exhibited a higher track length (Figure S3D), which could impact the number and duration of interactions. To control for this, we excluded any contact lasting less than one-second, and only quantified active exploration interactions. Together, these findings show that Lhx6^cKO^ animals engage in fewer social interactions.

**Figure 6.**
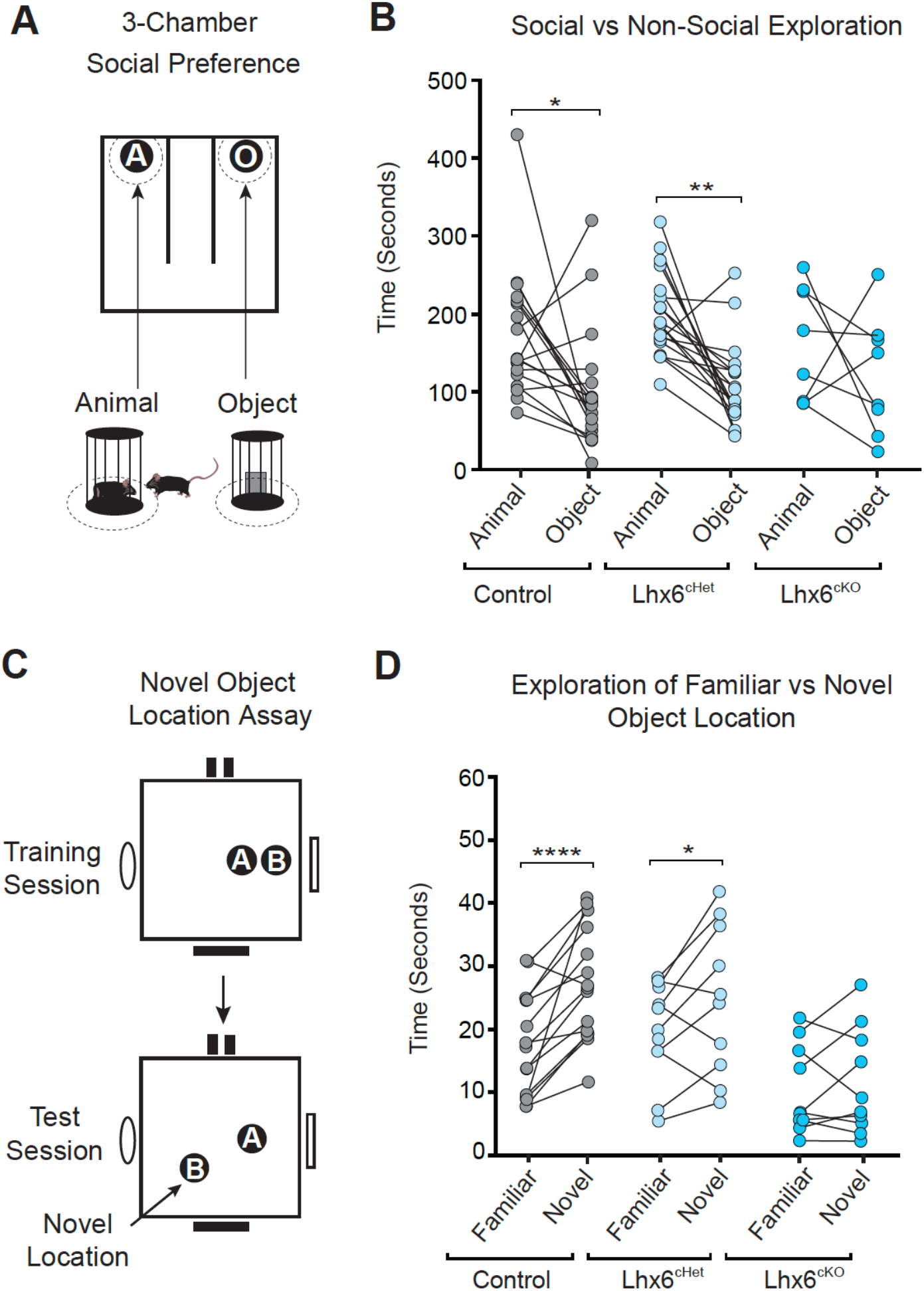
Embryonic loss of *Mef2c* from MGE-INs leads to changes in social and cognitive behavior. **(A)** Schematic of the 3-Chamber Social Preference assay, where animals explored chambers containing an Animal (novel conspecific mouse) or Object (binder clip) stimulus. Exploration time of the two stimuli was compared for each animal. **(B)** While only Control and Lhx6^cHet^ animals spent significantly more time with the social stimulus (Control n=18, Lhx6^cHet^ n=19, Lhx6^cKO^ n=8). Two-Way Repeated Measures ANOVA: stimulus only and stimulus*group interaction; paired measurements for each animal connected by line. **(C)** Schematic of the Novel Object Location Assay, where animals were exposed to two identical objects in a training session, followed by test session where one object was moved to a novel location within the arena. Exploration time of the two objects during the test session was compared for each animal. **(D)** Only control and Lhx6^cHet^ groups spent more time exploring the object that was moved to a novel location (Control n=15, Lhx6^cHet^ n=10, Lhx6^cKO^ n=10). For plots B and D, Two-way repeated measures ANOVA with Bonferroni post hoc comparison. * p < 0.05, ** p < 0.01, *** p< 0.001, ****p<0.0001. Data are presented as mean ± SEM. Detailed statistics in Table S4.

### Early removal of *Mef2c* from MGE-INs reduces exploration of novel objects

One typical behavioral feature of ASD is an increased preference for sameness or reduced exploration of novel changes to their environment (46). Accordingly, a previous study showed that knockdown of *Mef2c* from neuronal progenitors (Nestin-Cre:*Mef2c*-Flox) reduced exploration of novel environments (36). However, animals with removal of *Mef2c* from cortical excitatory neurons do not show such phenotypes (38), suggesting that novelty deficits arise from severe impairment of inhibitory activity. We used two novelty preference assays to probe whether animals can detect changes in object positioning (Novel Object Location) (Figure 6C-D) or object identity (Novel Object Recognition) (Figure S3E-F) within an open field arena. In both assays, animals are exposed to an initial set of objects and later timed on their exploration of the familiar versus novel object. Within the novel object location assay, all groups except Lhx6^cKO^ animals spent more time exploring the object in the new location (Figure 6C-D). Of note, Lhx6^cKO^ animals had lower overall exploration times, which may reflect the hyperlocomotion phenotype observed in the open field assay (Figure 4B). To control for this, we only included animals that reached a minimum exploration time for each object during the training session. Within the novel object recognition assay, none of the groups spent more time exploring the novel object (Figure S3E-F). Our findings of reduced spatial novelty recognition are consistent with other studies in which PV-IN development is impaired, including the Cantnap2 KO model and a Shank3 model with gene removal selectively in PV-INs (47, 48).

## Discussion

MEF2C is a transcription factor widely associated with various NDDs (2, 32, 49-52). MEF2C is among the highest ASD risk genes (51) and is strongly linked to schizophrenia (32), and intellectual disability (52). Here we employ genetic, electrophysiological, and behavioral approaches in mice to explore how cell-type specific loss of function of *Mef2c* at different ages leads to synaptic and neural circuit changes capable of recapitulating NDD-associated behaviors. We show that early, but not late, loss of *Mef2c* from PV-INs reduces their survival and reduces the excitatory drive onto remaining PV-INs. We also show that early *Mef2c* removal in Lhx6^cKO^ animals alters *in vivo* patterns of cortical activity suggestive of reduced inhibition and causes multiple behavioral phenotypes that are reminiscent of typical characteristics of ASD and other NDDs in humans. Taken together, our findings underscore the importance of *Mef2c* as a critical regulator of MGE-IN development, particularly PV-INs, and demonstrate how impaired interneuron development can lead to long-lasting changes in E/I synaptic balance, which likely underlies abnormal *in vivo* cortical activity, and contributes to various NDD behavioral phenotypes. Our findings are in agreement with previous work by Pai et al, which demonstrated that loss of Maf and Mafb transcription factors, which are shown to likely be upstream of *Mef2c*, leads to a reduction in MGE-lineage cells at age P35, with no change in the density of SST^+^ and a reduction of PV^+^ cells (53).

MEF2C loss-of-function mutations in humans are likely to lead to reduced inhibition given that *Mef2c* haploinsufficiency patients often develop seizures (6, 49, 50). Accordingly, mice lacking a copy of *Mef2c* have increased excitatory to inhibitory neurotransmission, suggesting reduced inhibition (7). We show that Lhx6^cKO^ animals have abnormal patterns of *in vivo* cortical activity consistent with reduced inhibition. Lhx6^cKO^ animals, but not PV^cKO^ animals, exhibit a change in broadband LFP power consistent with a decrease in the ratio of E/I conductance (34), a prevailing concept that unites the varied cellular dysfunctions that may give rise to pathophysiology present in NDDs (54). Lhx6^cKO^ animals may have reduced seizure threshold given the reduced E/I balance; however, we did not observe seizures in Lhx6^cKO^ animals. To further explore this, we plan to perform a study of cortical hyperactivity and seizure susceptibility in future studies.

MEF2C loss-of-function mutations in humans result in various NDD symptoms, including poor eye contact, absence of speech, intellectual disability, stereotypic movements, and hyperactivity (2, 4, 6). Accordingly, *Mef2c* haploinsufficiency mouse models exhibit NDD-relevant behavioral phenotypes such as impaired social interaction, hyperactivity, and cognitive deficits (1, 7, 55). Studies investigating the cell-specific role of *Mef2c* in PYR or microglia, have recapitulated some, but not all of the behavioral phenotypes observed in the mouse haploinsufficiency models (1, 3). We observed that early disruption of *Mef2c* in MGE-IN progenitors (Lhx6-Cre) resulted in hyperactivity, paw-clasping, reduced sociability, and differences in cognitive functioning. PV^cKO^ animals from our study and PV^cHet^ animals from a previous study (1) had no behavioral phenotypes, suggesting that MEF2C function in mature PV-INs is not critical for *Mef2c* pathophysiology. While at adult ages MEF2C is mostly expressed in PV-INs, it is possible that behavioral phenotypes observed in Lhx6^cKO^ are at least in part due to dysfunction of SST-INs. Furthermore, while our study focused on cortical interneurons, it is possible that Cre-dependent removal of *Mef2c* in non-neocortical regions may occur and contribute to the behavioral phenotypes observed.

Our results further show that *Mef2c* has a dosage-dependent effect. Loss of one copy of *Mef2c* in Lhx6^cHet^ mice leads to reduced PV-expression in PV-lineage INs, as well as reduced excitatory drive onto PV-INs to levels that are intermediary between Lhx6^cKO^ and control animals. Interestingly, this milder PV-IN phenotype in Lhx6^cHet^ mice did not lead to significant changes in in vivo patterns of network activity or changes in most of the tested NDD-associated behavioral phenotypes. In contrast, MEF2C haploinsufficiency both in humans and in mice appears to be sufficient to cause pathological behavior. It is likely that the compounded effects of loss of one copy of *Mef2c* in both inhibitory and excitatory neurons underlie the difference between the cell type specific loss of one copy of *Mef2c* and the haploinsufficiency model. Nevertheless, our findings show that *Mef2c* plays an essential role in the development of PV-INs at an early age, and that loss of *Mef2c* in cortical inhibitory neurons causes behaviors potentially relevant to multiple neurodevelopmental disorders.

## Acknowledgements

This work was supported by a Ruth L. Kirschstein Predoctoral Fellowship F31HD101360 to CW; a NARSAD Young Investigator Award, a Simons Bridge to Independence Award, a Whitehall Award, and National Institutes of Health (NIH) grants R01EY034617, R21MH133097, R01EY034310 to RBB; DA051608 to LS; NIH grants R01 NS113600 and R01 MH125772 to PEC. We thank Dr. Maria Gulinello for her guidance and support on rodent behavioral phenotyping.

## Competing interest statement

The authors report no biomedical financial interests or conflicts of interest.

Data in this paper are from a thesis to be submitted in partial fulfillment of the requirements for the Degree of Doctor of Philosophy in the Biomedical Sciences, Albert Einstein College of Medicine.

## Materials and Methods

### Animals

All mice were handled and maintained according to the regulations of the Institutional Animal Care and Use Committee at Albert Einstein College of Medicine. Lhx6-Cre (026555, Jackson Laboratory) and *Mef2c*-Floxed (025556, Jackson Laboratory) transgenic mouse lines were gifts of Dr. Gordon Fishell (Harvard Medical School, Broad Institute, USA). PV-Cre (017320, Jackson Laboratory) and SST-Cre (013044, Jackson Laboratory) mouse lines were obtained from Jackson Laboratory. The 5HT3aR-Cre transgenic mouse line was a gift from Dr. Nathaniel Heintz (The Rockefeller University). Mice were bred on-site to generate the experimental groups. Histological experiments were performed using Cre-positive animals obtained from crosses with a Cre-dependent fluorescent reporter, either RCE (010701, Jackson Laboratory) or Ai9 (007909, Jackson Laboratory) mouse lines. For all electrophysiology and behavioral experiments, the control group consisted of Cre-negative, *Mef2c*-floxed littermates (*Mef2c*F/F or *Mef2c*F/^+^).

### Histology

Histological characterizations were performed in the primary visual cortex (V1), the anterior cingulate cortex (ACC) and dentate gyrus of animals aged postnatal day 25 (P25) to P30. All image analysis was blinded to genotype. For quantification of MEF2C expression among interneuron subtypes, we compared the percentage of fate-mapped cells with MEF2C immunolabeling (PV-, SST-, or 5HT3A-Cre mice crossed to the RCE reporter mouse line). Total number of cells immunolabeled PV^+^ or SST^+^, and fate-mapped MGE-INs were normalized by optical area. MGE-INs were fate-mapped through crossing Lhx6-Cre mice to the Ai9 reporter mouse line, with and without floxed *Mef2c* alleles. Detailed protocols and reagent references are available in the Supplemental Methods section. Statistical details are available in tables S1-S2.

### Electrophysiology in acute brain slices

Whole-cell patch-clamp recordings were performed in PV-INs and pyramidal cells (PYRs) in V1 using a Multiclamp 700A amplifier (Molecular Devices). To identify PV-INs, pAAV-S5E2-C1V1-eYFP was injected in V1 and YFP-positive neurons were then visually patched (8). We confirmed that virally labeled PV-INs in control and mutant animals show typical fast spiking in response to depolarizing current injection (8). PYRs were identified using the soma shape. Recorded cells were filled with biocytin 4% and post hoc labeling with streptavidin-conjugated Alexa-647 was performed to confirm cell identity.

Voltage clamp recordings of Spontaneous Excitatory Postsynaptic Currents (sEPSCs) were recorded at Vholding = -60 mV in presence of GABAA (picrotoxin, 100 μM) and GABAB (CGP55845 hydrochloride, 3 μM) receptor antagonists. Spontaneous Inhibitory Postsynaptic Currents (sIPSCs) were recorded at Vholding = ^+^10 mV in the presence of AMPA (NBQX, 10 μM) and NMDA receptor (D-APV, 50 μM) antagonists. Electrophysiological data were acquired at 5 kHz, filtered at 2.4 kHz, and analyzed using custom-made software for IgorPro (Wavemetrics Inc.). Detailed solution preparations are available in the Supplemental Methods section. Statistical details are available in Table S3.

### Electrophysiology in awake head-fixed animals

In vivo electrophysiology recordings were performed in adult (4-8 months old) *Mef2c* mutants and littermate controls of both sexes. Recordings were conducted in non-anesthetized animals that were head-fixed and habituated to walk on a self-paced treadmill. Craniotomies were opened above ACC and V1 for the acute insertion of high-density silicone probes. Local field potential (LFP) analyses were performed using MATLAB-based scripts made available by the lab of György Buzsáki (New York University, https://github.com/buzsakilab/buzcode). In order to control for arousal state, all LFP power analyses were made using data collected during locomotion periods (see supplemental methods). Relative power in the specified frequency bands was measured as a ratio between power in those bands and the total power. Power spectra were normalized to total power for visualization purposes. To compute the slope of the aperiodic component of the LFP power spectrum, we use robust linear regression (MATLAB robustfit.m) to find the slope of the line best fit over the frequency range 0.5–100Hz. Statistical details are available in Table S4.

### Electrophysiology in awake head-fixed animals

Animals that underwent behavioral testing were group-housed and kept on a 14-hour light cycle with access to food and water ad libitum. Behavioral assays were performed during the light phase. The experimenter was blinded to the genotype of each test subject and the testing order was counterbalanced by experimental group. Mice were tested between 3-6 months of age. Male and female mice were pooled within each experimental group. Prior to each battery of behavior assays, animals were habituated to handling. Detailed protocols for each assay are available in the Supplemental Methods section. Statistical details are available in Tables S5-S7.

## Figures

**Figure S1.**
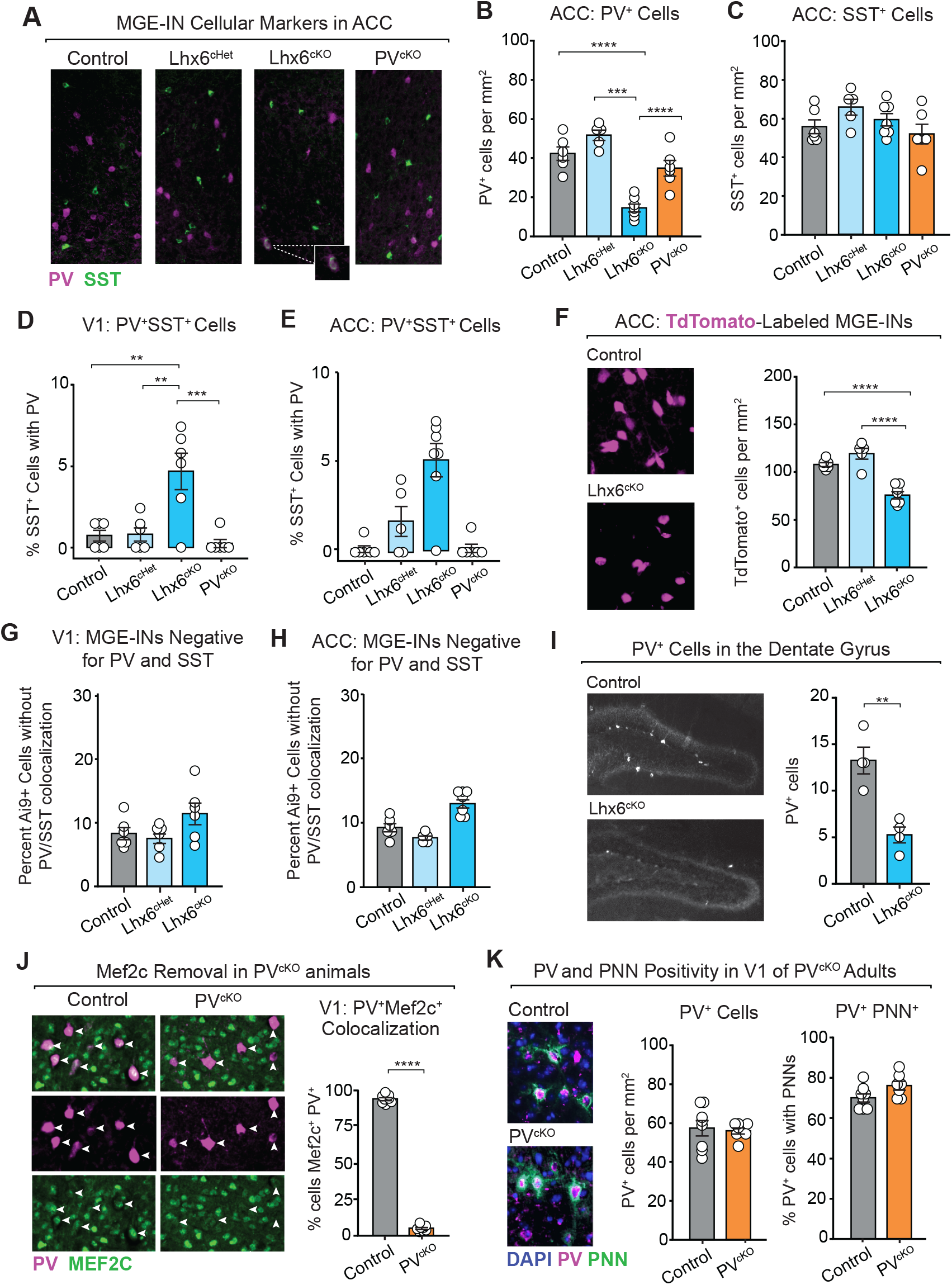
Embryonic removal of *Mef2c* from MGE-progenitors leads to a reduction in PV-INs, while PV^+^ IN number remains unchanged in adult animals with postnatal removal of *Mef2c* from MGE-INs. **(A)** MGE-IN immunopositivity for PV (magenta) and SST (green) in ACC. **(B)** PV^+^ cells normalized by area (cells per mm^2^) (control n=6, Lhx6^cHet^ n=5, Lhx6^cKO^ n=7, PV^cKO^ n=6). **(C)** SST^+^ cells in ACC normalized by area (cells per mm^2^) (control n=6, Lhx6^cHet^ n=5, Lhx6^cKO^ n=7, PV^cKO^ n=6). **(D**,**E)** Percentage of SST^+^ cells co-labelled with PV in V1 (control n=6, Lhx6^cHet^ n=6, Lhx6^cKO^ n=6, PV^cKO^ n=6) and ACC (control n=7, Lhx6^cHet^ n=5, Lhx6^cKO^ n=7, PV^cKO^ n=6). **(F)** TdTomato-labeling of MGE-INs was quantified and normalized by area (cells per mm^2^) (control n=6, Lhx6^cHet^ n=5, Lhx6^cKO^ n=7). **(G, H)** The percentage of PV/SST-negative MGE-INs out of the total number of MGE-INs (Cre-dependent labeling of Lhx6^+^ cells with TdTomato) in V1 (control n=6, Lhx6^cHet^ n=6, Lhx6^cKO^ n=6) and ACC (control n=6, Lhx6^cHet^ n=5, Lhx6^cKO^ n=7). **(I)** PV^+^ cells within the Dentate Gyrus compared between Lhx6^cKO^ animals (n=4) and littermate controls (n=4). (J) Confirmation of *Mef2c* (green) removal from PV^+^ cells (magenta) in 4-month-old PV^cKO^ animals (n=8) compared to littermate controls littermate controls (n=8). **(K, left)** PV^+^ cells within V1 normalized by area (cells per mm^2^) in 4-month-old PV^cKO^ animals (n=8) and littermate controls (n=7). **(K, right)** Percentage of PV^+^ cells within PNNS compared between 4-month-old PV^cKO^ animals (n=8) and littermate controls (n=7). (B-H) One-way ANOVA with Tukey HSD post-test. (I-K) Student’s t-test. *p<0.05, **p<0.01, ***p<0.001, **** p<0.0001. Each point corresponds to values from a single 10x field of view. Data are presented as mean ± SEM. For all groups n=6 (3 animals, 2-3 sections per animal). Detailed statistics in Table S2.

**Figure S2.**
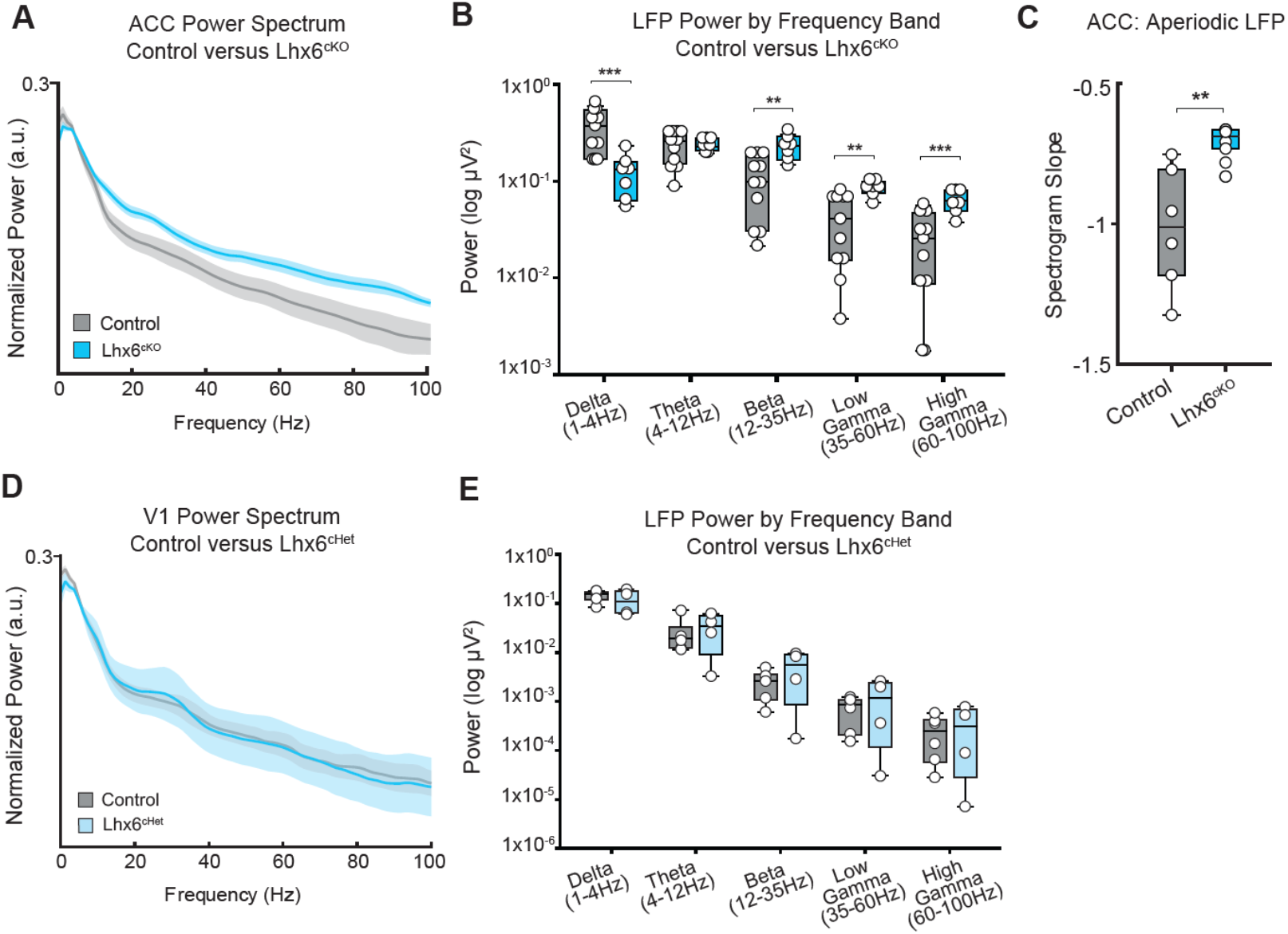
Embryonic loss of *Mef2c* in MGE-INs alters LFP power in ACC, Knockdown of *Mef2c* is not sufficient to alter LFP. **(A-B)** *In vivo* electrophysiology recordings from anterior cingulate cortex (ACC) performed in awake Lhx6^cKO^ mutants and littermate controls. **(A)** Population-averaged normalized ACC power spectra during locomotion periods for control (gray, n=6 session, x mice) and Lhx6^cKO^ animals (blue, n=4 sessions). **(B)** ACC relative LFP power compared between control (n=11 sessions) and Lhx6^cKO^ animals (n=7 sessions) across frequency bands during periods of locomotion. **(D-E)** *In vivo* electrophysiology recordings from primary visual cortex (V1) performed in awake Lhx6^cHet^ mutants (n=7 sessions) and littermate controls (n=7 sessions). **(D)** Population-averaged normalized V1 power spectra during locomotion periods for control (gray, n=4 sessions) and Lhx6^cHet^ animals (blue, n=4 sessions). **(E)** V1 LFP power compared between control (n=6 sessions) and Lhx6^cHet^ animals (n=4 sessions) across frequencies bands during periods of locomotion. Student’s t-test: * p < 0.05, ** p < 0.01, *** p< 0.001, ****p<0.0001. Each point corresponds to a single recording session. Data are presented as mean ± SEM. Detailed statistics in Table S4.

**Figure S3.**
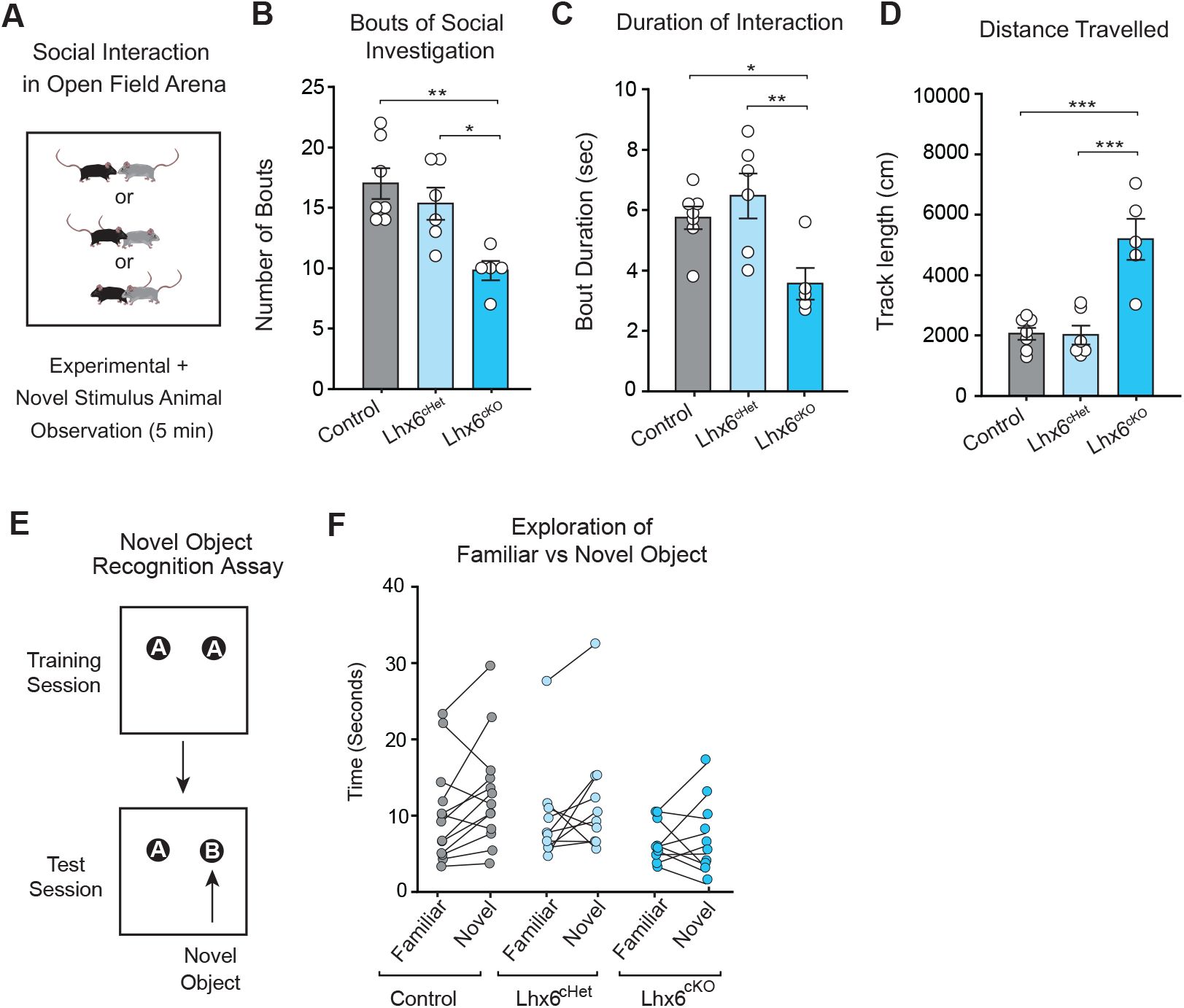
Additional behavioral assays. **(A-D)** Interaction between experimental animals and novel conspecific mice was scored for a five-minute period. Interactions were scored manually and were defined as proximal sniffing that was either mutual between the two animals, or sniffing carried out by a single animal toward the other (control n=13, Lhx6^cHet^ n=10, Lhx6^cKO^ n=10). **(B)** The number of social investigation bouts was lower in Lhx6^cKO^ animals, compared to control and Lhx6^cHet^ animals. **(C)** The duration of social investigation bouts was shorter in Lhx6^cKO^ animals. **(D)** Lhx6^cKO^ animals travelled a longer distance during the social interaction assay. **(E)** Schematic of the Novel Object Recognition assay in which the animal was exposed to two identical objects in a training session, followed by a test session where a novel object was introduced. **(F)** There was no significant difference in relative object exploration times within any of the groups (control n=13, Lhx6^cHet^ n=10, Lhx6^cKO^ n=10). For plots B-D, One-way ANOVA with Tukey HSD post hoc comparison. For plot F, Two-way repeated measures ANOVA with Bonferroni post hoc comparison. * p < 0.05, ** p < 0.01, *** p< 0.001, ****p<0.0001. Data are presented as mean ± SEM. Detailed statistics in Table S7.

## Supplemental Methods

### Histology

Animals were transcardially perfused with 4% Paraformaldehyde (PFA) in 1X Phosphate Buffered Saline (PBS), followed by a 1-hour tissue post-fixation at 4°C. Brains were prepared for cryosectioning with consecutive overnight incubations in 15% and 30% sucrose solutions in 1X PBS, followed by embedding in O.C.T. Compound embedding media (Fisher Scientific, Catalog number 23-730-571) on dry ice. Coronal sections were collected at a thickness of 20μn using a CryoStar™ NX70 cryostat (Epredia) onto SuperFrost™ plus microscope slides (Fisher Scientific, Catalog number 12-550-15). Slides were incubated for 1 hour at room temperature with a blocking solution containing 1.5% Normal Donkey Serum, 0.1% Triton-X-100 blocking solution prepared in 1X PBS. Primary and secondary antibodies were diluted using the blocking solution and all washes were carried out using 1X PBS. Slides were incubated with primary antibody for 48 hours at 4°C and secondary antibody for 1 hour at room temperature. During the primary incubation, tissues were either stained for MEF2C with rabbit Anti-MEF2C (1:300, Proteintech, Catalog number 10056-1-AP,) or co-stained with Mouse Anti-Parvalbumin (1:1000; Sigma-Aldrich, P3088), Guinea Pig Anti-Parvalbumin (1:500; P3088, SYSY Antibodies, Catalog number 195 308), and Rabbit Anti-Somatostatin (1:1000, Peninsula Laboratories, Catalog number T-4103,). Within the same primary antibody solution, the Cre-dependent TdTomato signal (Ai9) was amplified with Rat Anti-RFP (1:250; Chromotek, Catalog number 5F8). During the secondary incubation step, slides were incubated with Alexa Fluor™-conjugated anti-rabbit (1:500, Invitrogen, Catalog numbers A-21206, A-21207, A-31573), anti-rat (1:500, Invitrogen, Catalog numbers A-21208, A48271, A78947), anti-mouse antibodies (1:500, Invitrogen, Catalog numbers A-21202, A-21203, A-31571), and anti-guinea pig antibody (1:500, Jackson ImmunoResearch, Catalog number 706-605-148) (all raised in Donkey). Perineuronal nets were labeled by applying a biotinylated Wisteria Floribunda Lectin during the primary antibody incubation step (1:500; Vector Laboratories, Catalog number B-1355,) and an Alexa Fluor™-conjugated streptavidin during the secondary incubation step (1:500, Invitrogen, Catalog number S-11223). Slides were coverslipped with mounting media containing DAPI nuclear stain (ProLong™ Gold, Invitrogen, Catalog number P36935).

Slides were imaged on a Zeiss AXIO Imager.A1 microscope using a 10X or 20X objective. Analyses were conducted in the primary visual cortex (V1) and anterior cingulate cortex (ACC). To minimize counting bias, we compared sections of equivalent bregma positions, defined according to the Mouse Brain atlas (1).

### Electrophysiology in acute brain slices

Animals were anesthetized with isoflurane and euthanized in accordance with institutional regulations. Acute coronal slices (300 μm thick) were prepared from 4-9 month old mice using a VT1200s Microslicer (Leica Microsystems Co.) in a solution containing (in mM): 93 N-methyl-D-glucamine (NMDG), 2.5 KCl, 1.25 NaH2PO4, 30 NaHCO3, 20 HEPES, 25 D-glucose, 2 Thiourea, 5 Na-Ascorbate, 3 Na-Pyruvate, 0.5 CaCl2, 10 MgCl2, 93 HCl. Slices were then transferred to 32°C extracellular artificial cerebrospinal fluid (ACSF) solution, containing (in mM): 124 NaCl, 2.5 KCl, 26 NaHCO3, 1 NaH2PO4, 2.5 CaCl2, 1.3 MgSO4 and 10 D-glucose, for 30 min and then kept at room temperature for at least 40 min before electrophysiology recordings. All solutions were equilibrated with 95% O2 and 5% CO2 (pH 7.4).

Whole-cell patch-clamp recordings using a Multiclamp 700A amplifier (Molecular Devices) were performed in parvalbumin-expressing interneurons (PV-INs) and pyramidal cells (PYRs) in V1. To identify PV-INs, AAV-S5E2C1V1-YFP was injected in V1 and YFP-positive neurons were then visually patched (2). We confirmed that virally labeled PV-INs show typical fast spiking in response to depolarizing current injection (2). PYRs were identified using the soma shape. Recorded cells were filled with biocytin 4% and post hoc labeling with streptavidin-conjugated Alexa-647 was performed to confirm cell identity.

Inhibitory postsynaptic currents (IPSCs) were monitored in voltage clamp mode using patch-type pipette electrodes (∼3-4 MΩ) containing (in mM): 131 cesium gluconate, 8 NaCl, 1 CaCl2, 10 EGTA, 10 D-glucose and 10 HEPES, pH 7.2 (285-290 mOsm). Spontaneous IPSCs (sIPSCs) were recorded at V_holding_ = ^+^10 mV in the presence of AMPA (NBQX, 10 μM) and NMDA receptor (D-APV, 50 μM) antagonists. Voltage clamp recording of excitatory postsynaptic currents (EPSCs) and current clamp recordings were performed using K^+^-based internal solution containing (in mM): 135 KMeSO4, 5 KCl, 1 CaCl2, 5 NaOH, 10 HEPES, 5 MgATP, 0.4 Na3GTP, 5 EGTA and 10 D-glucose, pH 7.2 (288-292 mOsm). Spontaneous EPSCs (sEPSCs) were recorded at V_holding_ = -60 mV in presence of GABAA (picrotoxin, 100 μM) and GABAB (CGP55845 hydrochloride, 3 μM) receptor antagonists. All recordings were performed at 28 ± 1°C in a submersion-type recording chamber perfused at 2 ml/min with ACSF. Series resistance (∼6-21 MΩ) was monitored throughout all experiments with a −5 mV, 80 ms voltage step, and cells that exhibited a significant change in series resistance (> 20%) were excluded from analysis. Electrophysiological data were acquired at 5 kHz, filtered at 2.4 kHz, and analyzed using custom-made software for IgorPro (Wavemetrics Inc.). Reagents were bath applied following dilution into ACSF from stock solutions stored at −20°C prepared in water or DMSO, depending on the manufacturer’s recommendation.

### Electrophysiology in awake head-fixed animals

Headpost Surgery and Treadmill Training—*In vivo* electrophysiology recordings were performed in adult (2-12 months old) Mef2c mutants and littermate controls of both sexes. Recordings were conducted in non-anesthetized animals that were head-fixed and able to walk on a self-paced treadmill. Mice were initially handled for 5–10 min/day for 5 days prior to a headpost surgery. On the day of the surgery, the mouse was anesthetized with isoflurane (3%–5% induction, 1.5% maintenance) 10 min after injection of a systemic analgesic (meloxicam, 5 mg per kg of body weight) and placed in a stereotaxic frame. Mice were kept at 37°C at all times using a feedback-controlled heating pad. Pressure points and incision sites were injected with lidocaine (2%), and eyes were protected from desiccation using artificial tear ointment. The skin above the skull was incised and a custom-made 3D-printed headpost was implanted on the skull using Vetbond (3M), leaving a bilateral window of skull uncovered over the anterior cingulate cortex and primary visual cortex. A recording chamber was built using dental cement (Ortho-Jet, Lang). Analgesics were given immediately after the surgery and on the two following days to aid recovery. Mice were returned to their home cage and allowed to recover for 3–5 days following implant surgery before beginning treadmill training.

Once recovered from the surgery, mice were trained for head-fixation to bars placed near the front of the treadmill apparatus. Headposts were custom designed to mimic the natural head angle of the running mouse. The treadmill was 3D-printed in plastic with dimensions 15cm long by 6cm wide (Janelia Open Science Laboratory Tools). Both treadmill and headpost bars were mounted on an air table to minimize vibrations and electrical noise during the electrophysiology session. A programmable magnetic angle sensor (Digikey) was attached to the rear axel of the treadmill for continuous monitoring of locomotion. During training, mice were headposted for increasing intervals on each successive day. If signs of anxiety or distress were noted, the mouse was removed from the headpost and the training interval was not lengthened on the next day. Mice were trained on the wheel for up to 7 days or until they exhibited robust bouts of running activity during each session. Mice that continued to exhibit signs of distress were not used for awake electrophysiology sessions.

Craniotomy and Electrophysiology Recordings—Two circular craniotomies (diameter 3 mm) were performed above ACC and V1, and the recording chamber was covered with a silicone adhesive (Kwik-Sil). The mouse was returned to its home cage to recover from anesthesia. On the day of the recording, the mouse was headfixed onto the treadmill and the silicone adhesive was removed. Two high-density silicon electrodes (H3 Cambridge Neuronexus) were acutely inserted into both ACC and V1. Each probe had 1 shank with 64 electrode contacts (area of each contact 0.02 μm^2^, probe length 1.275 mm, and 86 μm at its widest point and tapered to a tip). Each electrode was connected to a headstage (IntanTechnologies, RHD2000 64-channel Amplifier Board with two RHD2164 amplifier chips) and the headstage was connected to an Intan RHD2000 Evaluation Board, which sampled each signal at a rate of 20 kHz per channel. To ensure synchrony between physiological signals and locomotion, the linear angular detector signals were measured in tandem with electrophysiological signals by the same IntanRHD2000 Evaluation Board. Data were recorded using Intan RHX acquisition software and saved as binary files.

### Behavioral assays

Animals were group housed and kept on a 14-hour light cycle (lights on at 6AM) with access to food and water ad libitum. Behavioral assays were performed during the light phase. The experimenter was blinded to the genotype of each test-subject and testing order was counterbalanced by experimental group. Mice were tested between 3-6 months of age. Male and female mice were pooled within each experimental group. Prior to each battery of behavior assays, animals were habituated to handling by the experimenter for three 5-minute sessions.

#### Paw Clasping

Paw clasping was measured as the percentage of time the animal spent with any reflexive limb clasping during a two-minute tail suspension assay. During the tail suspension assay, the experimenter held the animal one centimeter above the base of the tail with their thumb, index and middle fingers. The assay was performed over a lab benchtop with the experimenter resting their forearm a box such that the animal was 30cm from both the benchtop and the wall to ensure both stability and consistency. The assay was video recorded and scored post-hoc with a stopwatch at 0.5x speed. Periods of clasping included retraction of one or more limbs toward the body, which often included the crossing of front paws, hind paws, or all paws. Given the distress inherent to this test, tail-suspension was performed after all other behavior assays were completed. The assay was terminated early for any animal exhibiting signs of extreme distress, such as vocalization.

#### Open Field Test (OFT)

Each mouse was run in the OFT after completing three 5-minute handling sessions conducted one day prior to testing in order to reduce the amount of stress when transferring the animal from the home cage to the open field arena. In this test, each mouse was allowed to freely explore the open field (square arena: 40.5 cm length x 40.5 cm width x 30.5 cm height) for 12 minutes. In a subset of animals, measurements of tracklength, jumping and grooming activity were measured using the Behavior Sequencer apparatus from Biobserve (Bonn, Germany), which employs a pattern recognition algorithm to classify the activity of the animal based on vibrational and postural measurements recorded from combination of piezoelectric and infrared sensors, and video tracking.

#### Elevated Plus Maze (EPM)

The EPM apparatus (Stoelting Co., Wood Dale, IL; Product ID: 60240) was a plus-shaped maze with 4 equally-sized rectangular arms (50 x 10 cm) united by a square-shaped center (10 x 10 cm), elevated 50 cm above the floor. It consisted of two closed arms enclosed on three sides by opaque walls (40 cm high), and two open arms. For testing in the EPM each animal was placed in the center and allowed to freely explore the maze for 5 minutes. Measurements of tracklength and duration in each arm was calculated with an automated video-tracking system (Viewer, Biobserve, Germany). The open and closed arm zones were defined in the software as beginning where a mouse’s body could be fully in a given arm.

#### Y-Maze

The Y-Maze apparatus (Stoelting Co., Wood Dale, IL; Product ID: 60180) was a plus-shaped maze with 3 equally-sized rectangular arms (35 x 5 cm) united by a triangle-shaped center (each side 5 cm). Walls (10cm height) were present on arms of the maze and joined together to enclose the maze center. For testing in the Y-maze each animal was placed in the center and allowed to freely explore the maze for 5 minutes. Measurements of tracklength and duration in each arm were calculated with an automated video-tracking system (Viewer, Biobserve, Germany). The number of times the animal consecutively alternated through all three arms (Arms: A, B, C) was scored from videos post-hoc by an observer blinded to animal genotype. Entires that resulted in one of the following three sequences were considered “spontaneous alternations”: A-B-C, A-C-B, B-A-C, B-C-A, C-A-B, C-B-A. The percentage of spontaneous alternations was calculated as: [((number of spontaneous alternations)/(total arm entries))*100]. The first two entries were not counted toward the total number of entries, as three entries were required to make a full sequence. Entries were divided into full and partial, where animals reached the end of the maze in full entries. Entries were considered partial when the animal was within a zone covering the two middle quarters of the maze arm before returning to the center of the maze. Movements into the first quarter of the maze from the center of the maze were not considered entries, as these often included instances of the animal turning in the center.

#### Social Preference (SP)

The SP apparatus consisted of a custom 3-chamber arena, which was a modification of the 40.5 cm length x 40.5 cm width x 30.5 cm height open field arena. The arena was divided into three chambers by inserting two walls, which were the same height as the outer walls and extended 24.7 cm from the back wall of the arena. The walls were spaced such that each outer chamber was 15.25cm wide and the central chamber was 8cm wide. Each outer chamber contained a wire cup at the center of the back wall (8.5cm diameter) which housed either a stimulus animal or a non-social object. An automated video-tracking system (Viewer, Biobserve, Germany) was used to draw zones around the stimuli (13cm). Investigation duration for each stimulus was determined through tracking the nose of the test animal. For SP testing, each animal was placed in the center chamber and allowed to freely explore the arena for 10 minutes while wire cups were empty. After 10 minutes the animal was guided to the center chamber and the exit was blocked while the stimulus animal and non-social object were placed within the cups. Animals were then allowed to explore the full arena for 10 minutes. SP data are reported as the raw duration of test animals interacting with animal and object stimuli. Stimulus animals were novel sex- and age-matched conspecifics and the non-social object was a binder clip. All stimulus animals were habituated to the wire cup in three 10-minute sessions on the day prior to the SP assay.

#### Social Interaction

The assay was carried out in an open field arena with dimensions 40.5 cm length x 40.5 cm width x 30.5 cm height open field. Animals were allowed to explore the open field arena for 10 minutes prior to the introduction on a novel conspecific stimulus animal. Aged (12 months), senescent female conspecific stimulus animals were used to limit estrous cycle related variations in social exploration. For ease of identification, stimulus animals were on a BALB/c background (white fur) to contrast with the black coat color of experimental animals. Scoring of social interactions was carried out manually and an automated video-tracking system (Viewer, Biobserve, Germany) was used to measure track length for the experimental animal. Investigation duration was defined as proximal sniffing that was either mutual between the two animals or carried out by one animal toward the other. Sniffing was considered proximal sniffing if the nose of the animal was within roughly 1cm or in direct contact with the other animal. Bouts of interaction that lasted less than one second were excluded from analyses. All stimulus animals were habituated to handling by the experimenter the day prior to the SI assay to reduce stress.

#### Novel Object Location Test

A square-shaped arena (40.5 cm length x 40.5 cm width x 30.5 cm height) was set up with two novel and identical objects on the floor. The arena contained unique high-contrast visual cues to aid the animals in determining the object placement, and visually rich objects were used to encourage higher levels of object exploration. Mice were individually placed in the arena for a 5-minute training session in which mice were allowed to freely explore both objects. Animals were then removed from the arena for a twenty-minute retention interval during which one of the objects was moved 12 cm from its original position. Following the retention interval, animals were returned to the arena for a test session freely explore the two objects, for 5 minutes. Exploration of the objects was quantified by an experimenter and confirmed with computer tracking (Viewer, Biobserve). Mice that had a total object exploration time less than 3 seconds in training or testing were excluded.

#### Novel Object Recognition Test

A square-shaped arena (40.5 cm length x 40.5 cm width x 30.5 cm height) was set up with two novel and identical objects on the floor. During training, mice were allowed to freely explore the two identical objects for 4 minutes. For testing, one of the objects was replaced with another novel object (new object) but kept in the same position as the original object. Forty-five minutes after training, the testing session was performed in which each mouse was allowed to freely explore the old and new objects for 5 minutes. Novel object position (left or right) was randomized. Exploration of the objects was quantified by an experimenter and confirmed with computer tracking (Viewer, Biobserve). Mice that had a total object exploration time less than 3 seconds in training or testing were excluded.

## Supplemental Tables

**Table S1.** Comparison of Conditional *Mef2c* Removal on MGE-IN subtypes in V1. *(Corresponding to Figure 1)*

| Figure | Comparison | N | Test<sup>a</sup> | Comparison | p-value<sup>b</sup> | Level<sup>c</sup> |
| --- | --- | --- | --- | --- | --- | --- |
| 1B | Interneuron lineage-specific GFP expression stained with anti-Mef2c | All groups = 3 animals with 2-5 sections each<br>PV-Cre = 10<br>SST-Cre = 10<br>5HT3A-Cre = 10 | One-way ANOVA<br>Tukey HSD | All Groups<br>PV-Cre vs. SST-Cre<br>PV-Cre vs. 5HT3A-Cre<br>SST-Cre vs. 5HT3A-Cre | F(2,27)= 785.7, p<0.0001<br><0.0001<br><0.0001<br><0.0001 | ****<br>****<br>****<br>**** |
| 1E | PV+ Cells per mm<sup>2</sup> in V1 | All groups = 3 animals with 2 sections each<br>Control = 6<br>Lhx6<sup>chHet</sup> = 6<br>Lhx6<sup>ckO</sup> = 6<br>PV<sup>ckO</sup> = 6 | One-way ANOVA<br>Tukey HSD | All Groups<br>Control vs. Lhx6<sup>chHet</sup><br>Control vs. Lhx6<sup>ckO</sup><br>Control vs. PV<sup>ckO</sup><br>Lhx6<sup>chHet</sup> vs. Lhx6<sup>ckO</sup><br>PV<sup>ckO</sup> vs. Lhx6<sup>ckO</sup> | F(3,20)= 19.6, p<0.0001<br>0.0060<br><0.0001<br>0.4464<br>0.0128<br><0.0001 | ****<br>**<br>****<br>*ns*<br>*<br>**** |
| 1F | SST+ Cells per mm<sup>2</sup> in V1 | All groups = 3 animals with 1-2 sections each<br>Control = 6<br>Lhx6<sup>chHet</sup> = 6<br>Lhx6<sup>ckO</sup> = 6<br>PV<sup>ckO</sup> = 5 | One-way ANOVA | All Groups<br>Control vs. Lhx6<sup>chHet</sup><br>Control vs. Lhx6<sup>ckO</sup><br>Control vs. PV<sup>ckO</sup><br>Lhx6<sup>chHet</sup> vs. Lhx6<sup>ckO</sup><br>PV<sup>ckO</sup> vs. Lhx6<sup>ckO</sup> | F(3,20)= 26.0, p=0.0496<br>0.9998<br>0.2940<br>0.0940<br>0.3337<br>0.9102 | *<br>*ns*<br>*ns*<br>*ns*<br>*ns*<br>*ns* |
| 1H | TdTomato-Labeled MGE-INs (Lhx6-expressing cells) | All groups = 3 animals with 2 sections each<br>Control = 6<br>Lhx6<sup>chHet</sup> = 6<br>Lhx6<sup>ckO</sup> = 6 | One-way ANOVA<br>Tukey HSD | All Groups<br>Control vs. Lhx6<sup>chHet</sup><br>Control vs. Lhx6<sup>ckO</sup><br>Lhx6<sup>chHet</sup> vs. Lhx6<sup>ckO</sup> | F(2,15)= 19.6, p<0.0001<br>0.1627<br><0.0001<br>0.0021 | ****<br>*ns*<br>****<br>** |
| 1J | PV-INs with Perineuronal Nets | Control = 6<br>Lhx6<sup>chHet</sup> = 6<br>Lhx6<sup>ckO</sup> = 6<br>PV<sup>ckO</sup> = 6 | One-way ANOVA<br>Tukey HSD | All Groups<br>Control vs. Lhx6<sup>chHet</sup><br>Control vs. Lhx6<sup>ckO</sup><br>Control vs. PV<sup>ckO</sup><br>Lhx6<sup>chHet</sup> vs. Lhx6<sup>ckO</sup><br>PV<sup>ckO</sup> vs. Lhx6<sup>ckO</sup> | F(3,20)= 22.6, p<0.0001<br>0.9987<br><0.0001<br>0.6166<br><0.0001<br><0.0001 | <0.0001<br>*ns*<br>****<br>*ns*<br>****<br>**** |

<sup>a</sup>For significant multiple comparisons (ANOVA), a Tukey HSD post hoc test was applied for determining pairwise differences<sup>b</sup>alpha = 0.05<sup>c</sup>Notation for significance levels: \*p<0.05, \*\*p<0.01, \*\*\*p<0.001, \*\*\*\* p<0.0001, *ns* = not significant (p>0.05)

**Table S2.** Comparison of Conditional *Mef2c* Removal on MGE-IN subtypes in V1 and ACC. *(Corresponding to Figure S1)*

| Figure | Comparison | N | Test<sup>a</sup> | Comparison | p-value<sup>b</sup> | Level<sup>c</sup> |
| --- | --- | --- | --- | --- | --- | --- |
| S1B | PV+ Cells per mm<sup>2</sup> in ACC | Control = 6<br>Lhx6<sup>cHet</sup> = 5<br>Lhx6<sup>cKO</sup> = 7<br>PV<sup>cKO</sup> = 6<br>(3 animals, 1-3 sections) | One-way ANOVA<br>Tukey HSD | All Groups<br>Control vs. Lhx6<sup>cHet</sup><br>Control vs. Lhx6<sup>cKO</sup><br>Control vs. PV<sup>cKO</sup><br>Lhx6<sup>cHet</sup> vs. Lhx6<sup>cKO</sup><br>PV<sup>cKO</sup> vs. Lhx6<sup>cKO</sup> | F(3,20)= 27.0,<br>p<0.0001<br>0.2036<br><0.0001<br>0.3445<br>0.0006<br><0.0001 | ****<br>*ns*<br>****<br>*ns*<br>***<br>**** |
| S1C | SST+ Cells per mm<sup>2</sup> in ACC | Control = 6<br>Lhx6<sup>cHet</sup> = 5<br>Lhx6<sup>cKO</sup> = 7<br>PV<sup>cKO</sup> = 6<br>(3 animals, 1-3 sections) | One-way ANOVA | All Groups | F(3,20)= 1.9,<br>p=0.1584 | *ns* |
| S1D | Percent of SST+ cells that co-express PV in V1 | Control = 6<br>Lhx6<sup>cHet</sup> = 6<br>Lhx6<sup>cKO</sup> = 6<br>PV<sup>cKO</sup> = 6<br>(3 animals, 1-3 sections) | One-way ANOVA<br>Tukey HSD | All Groups<br>Control vs. Lhx6<sup>cHet</sup><br>Control vs. Lhx6<sup>cKO</sup><br>Control vs. PV<sup>cKO</sup><br>Lhx6<sup>cHet</sup> vs. Lhx6<sup>cKO</sup><br>PV<sup>cKO</sup> vs. Lhx6<sup>cKO</sup> | F(3,20)= 10.7,<br>p=0.0002<br>1.0000<br>0.0013<br>0.9476<br>0.0015<br>0.0004 | ***<br>*ns*<br>**<br>*ns*<br>**<br>*** |
| S1E | Percent of SST+ cells that co-express PV in ACC | Control = 7<br>Lhx6<sup>cHet</sup> = 5<br>Lhx6<sup>cKO</sup> = 7<br>PV<sup>cKO</sup> = 6<br>(3 animals, 1-3 sections) | One-way ANOVA<br>Tukey HSD | All Groups<br>Control vs. Lhx6<sup>cHet</sup><br>Control vs. Lhx6<sup>cKO</sup><br>Control vs. PV<sup>cKO</sup><br>Lhx6<sup>cHet</sup> vs. Lhx6<sup>cKO</sup><br>PV<sup>cKO</sup> vs. Lhx6<sup>cKO</sup> | F(3,20)= 13.1,<br>p<0.0001<br>0.4178<br>0.0002<br>1.0000<br>0.0113<br>0.0002 | ****<br>*ns*<br>***<br>*ns*<br>*<br>*** |
| S1F | TdTomato-Labeled MGE-INs (Lhx6-expressing cells) in ACC | Control = 6<br>Lhx6<sup>cHet</sup> = 5<br>Lhx6<sup>cKO</sup> = 7<br>(3 animals, 1-3 sections) | One-way ANOVA<br>Tukey HSD | All Groups<br>Control vs. Lhx6<sup>cHet</sup><br>Control vs. Lhx6<sup>cKO</sup><br>Lhx6<sup>cHet</sup> vs. Lhx6<sup>cKO</sup> | F(2,15)= 34.0,<br>p<0.0001<br>0.1570<br><0.0001<br><0.0001 | ****<br>*ns*<br>****<br>**** |

<sup>a</sup>For significant multiple comparisons (ANOVA), a Tukey HSD post hoc test was applied for determining pairwise differences<sup>b</sup>alpha = 0.05<sup>c</sup>Notation for significance levels: \*p<0.05, \*\*p<0.01, \*\*\*p<0.001, \*\*\*\* p<0.0001, *ns* = not significant (p>0.05)

**Table S2 Continued. Comparison of Conditional *Mef2c* Removal on MGE-IN subtypes in V1 and ACC** (Corresponding to Figure S1)
| Figure | Comparison | N | Test<sup>a</sup> | Comparison | p-value<sup>b</sup> | Level<sup>c</sup> |
| --- | --- | --- | --- | --- | --- | --- |
| S1G | PV and SST Negative MGE-INs in V1 | Control = 6<br>Lhx6<sup>cHet</sup> = 6<br>Lhx6<sup>cKO</sup> = 6<br>(3 animals, 1-3 sections) | One-way ANOVA | All Groups | F(2,15)= 22.2,<br>p=0.0830 | ns |
| S1H | PV and SST Negative MGE-INs in ACC | Control = 6<br>Lhx6<sup>cHet</sup> = 5<br>Lhx6<sup>cKO</sup> = 7<br>(3 animals, 1-3 sections) | One-way ANOVA<br>Tukey HSD | All Groups<br>Control vs. Lhx6<sup>cHet</sup><br>Control vs. Lhx6<sup>cKO</sup><br>Lhx6<sup>cHet</sup> vs. Lhx6<sup>cKO</sup> | F(2,15)= 22.2,<br>p<0.0001<br>0.1863<br>0.0009<br><0.0001 | ****<br>ns<br>***<br>**** |
| S1I | PV+ cells in Dentate Gyrus | Control = 4<br>Lhx6<sup>cKO</sup> = 4<br>(3 animals, 1-2 sections) | Student's t-test | Control vs. Lhx6<sup>cKO</sup> | 0.0052 | ** |
| S1J | Removal of *Mef2c* from PV+ cells at 4 months | Control = 8<br>PV<sup>cKO</sup> = 8<br>(2 animals, 1-3 sections) | Student's t-test | Control vs. PV<sup>cKO</sup> | <0.0001 | **** |
| S1K left | PV+ cells in V1 at 4 months | Control = 8<br>PV<sup>cKO</sup> = 7<br>(2 animals, 4 sections) | Student's t-test | Control vs. PV<sup>cKO</sup> | 0.7526 | ns |
| S1K right | Percent PV+ cells with PNNs at 4 months | Control = 8<br>PV<sup>cKO</sup> = 7<br>(2 animals, 3-4 sections) | Student's t-test | Control vs. PV<sup>cKO</sup> | 0.0621 | ns |

<sup>a</sup>For significant multiple comparisons (ANOVA), a Tukey HSD post hoc test was applied for determining pairwise differences<sup>b</sup>alpha = 0.05<sup>c</sup>Notation for significance levels: \*p<0.05, \*\*p<0.01, \*\*\*p<0.001, \*\*\*\* p<0.0001, ns = not significant (p>0.05)

**Table S3.** Comparison of Conditional *Mef2c* Removal on Cellular Function. *(Corresponding to Figure 2)*

| Figure | Comparison | N | Test<sup>a</sup> | Comparison | p-value<sup>b</sup> | Level<sup>c</sup> |
| --- | --- | --- | --- | --- | --- | --- |
| 2D | PV-IN sEPSC Frequency | Control = 8, 4 mice<br>Lhx6<sup>chHet</sup> = 6, 3 mice<br>Lhx6<sup>cKO</sup> = 6, 3 mice<br>PV<sup>cKO</sup> = 7, 2 mice | One-way ANOVA<br>Tukey HSD | All Groups<br>Control vs. Lhx6<sup>chHet</sup><br>Control vs. Lhx6<sup>cKO</sup><br>Control vs. PV<sup>cKO</sup><br>Lhx6<sup>chHet</sup> vs. Lhx6<sup>cKO</sup><br>PV<sup>cKO</sup> vs. Lhx6<sup>cKO</sup> | F(3,26)= 27.0, p < 0.0001<br>p < 0.01<br>p < 0.001<br>p = 0.82<br>p < 0.01<br>p < 0.0001 | ****<br>**<br>***<br>*ns*<br>**<br>**** |
| 2D | sEPSC Amplitude | Control = 8, 4 mice<br>Lhx6<sup>chHet</sup> = 6, 3 mice<br>Lhx6<sup>cKO</sup> = 6, 3 mice<br>PV<sup>cKO</sup> = 7, 2 mice | One-way ANOVA | All Groups | F(3,26)= 1.08, p = 0.38 | *ns* |
| 2F | Pyramidal Cell sIPSC Frequency | Control = 8, 5 mice<br>Lhx6<sup>chHet</sup> = 8, 4 mice<br>Lhx6<sup>cKO</sup> = 7, 3 mice | One-way ANOVA<br>Tukey HSD | All Groups<br>Control vs. Lhx6<sup>chHet</sup><br>Control vs. Lhx6<sup>cKO</sup><br>Lhx6<sup>chHet</sup> vs. Lhx6<sup>cKO</sup> | F(2,22)= 10.8, p < 0.001<br>< 0.05<br>< 0.001<br>0.17 | ***<br>*<br>***<br>*ns* |
| 2F | Pyramidal Cell sIPSC Amplitude | Control = 8, 5 mice<br>Lhx6<sup>chHet</sup> = 8, 4 mice<br>Lhx6<sup>cKO</sup> = 7, 3 mice | One-way ANOVA | All Groups | F(2,22)= 0.13, p = 0.88 | *ns* |

<sup>1</sup>For significant multiple comparisons (ANOVA), a Tukey HSD post hoc test was applied for determining pairwise differences<sup>2</sup>Statistically significant comparisons are highlighted in red text, alpha = 0.05<sup>3</sup>Notation for significance levels: \*p<0.05, \*\*p<0.01, \*\*\*p<0.001, \*\*\*\* p<0.0001, *ns* = not significant (p>0.05)

**Table S4.** Comparison of Conditional *Mef2c* Removal on Network Activity. *(Corresponding to Figure 3, Figure S2)*

| Figure | Comparison | N | Test | Comparison | p-value<sup>a</sup> | Level<sup>b</sup> |
| --- | --- | --- | --- | --- | --- | --- |
| 3D | V1 LFP power across frequency bands<br>Control versus Lhx6<sup>ckO</sup> | Control = 6<br>Lhx6<sup>ckO</sup> = 4 | Student's t-test | V1: 1-4 Hz Power | 0.0103 | * |
|  |  |  |  | V1: 4-12 Hz Power | 0.7772 | ns |
|  |  |  |  | V1: 12-35 Hz Power | 0.0066 | ** |
|  |  |  |  | V1: 35-60 Hz Power | 0.0048 | ** |
|  |  |  |  | V1: 60-100 Hz Power | 0.0180 | * |
| 3E | V1 LFP Aperiodic Slope<br>Control versus Lhx6<sup>ckO</sup> | Control = 7<br>Lhx6<sup>ckO</sup> = 7 | Student's t-test | Control versus Lhx6<sup>ckO</sup> | 0.0107 | * |
| 3G | V1 LFP power across frequency bands<br>Control versus PV<sup>ckO</sup> | Control = 4<br>PV<sup>ckO</sup> = 4 | Student's t-test | V1: 1-4 Hz Power | 0.5409 | ns |
|  |  |  |  | V1: 4-12 Hz Power | 0.9059 | ns |
|  |  |  |  | V1: 12-35 Hz Power | 0.6484 | ns |
|  |  |  |  | V1: 35-60 Hz Power | 0.8376 | ns |
|  |  |  |  | V1: 60-100 Hz Power | 0.3312 | ns |
| 3H | V1 LFP Aperiodic Slope<br>Control versus PV<sup>ckO</sup> | Control = 4<br>PV<sup>ckO</sup> = 4 | Student's t-test | Control versus PV<sup>ckO</sup> | 0.2188 | ns |
| S2B | ACC LFP power across frequency bands<br>Control versus Lhx6<sup>ckO</sup> | Control = 11<br>Lhx6<sup>ckO</sup> = 7 | Student's t-test | ACC: 1-4 Hz Power | 0.0009 | *** |
|  |  |  |  | ACC: 4-12 Hz Power | 0.8980 | ns |
|  |  |  |  | ACC: 12-35 Hz Power | 0.0021 | ** |
|  |  |  |  | ACC: 35-60 Hz Power | 0.0012 | ** |
|  |  |  |  | ACC: 60-100 Hz Power | 0.0007 | *** |
| S2C | ACC LFP Aperiodic Slope<br>Control versus Lhx6<sup>ckO</sup> | Control = 7<br>Lhx6<sup>ckO</sup> = 7 | Student's t-test | Control versus Lhx6<sup>ckO</sup> | 0.0040 | ** |
| S2E | V1 LFP power across frequency bands<br>Control versus Lhx6<sup>chHet</sup> | Control = 6<br>Lhx6<sup>chHet</sup> = 4 | Student's t-test | V1: 1-4 Hz Power | 0.4769 | ns |
|  |  |  |  | V1: 4-12 Hz Power | 0.6483 | ns |
|  |  |  |  | V1: 12-35 Hz Power | 0.3223 | ns |
|  |  |  |  | V1: 35-60 Hz Power | 0.4820 | ns |
|  |  |  |  | V1: 60-100 Hz Power | 0.6851 | ns |

<sup>a</sup>alpha = 0.05<sup>b</sup>Notation for significance levels: \*p<0.05, \*\*p<0.01, \*\*\*p<0.001, \*\*\*\* p<0.0001, ns = not significant (p>0.05)

**Table S4 Continued. Comparison of Conditional *Mef2c* Removal on Network Activity** (Corresponding to Figure 3, Figure S2)
| Figure | Comparison | N | Test | Comparison | p-value<sup>a</sup> | Level<sup>b</sup> |
| --- | --- | --- | --- | --- | --- | --- |
| S2C | ACC LFP Aperiodic Slope<br>Control versus Lhx6<sup>ckO</sup> | Control = 7<br>Lhx6<sup>ckO</sup> = 7 | Student's t-test | Control versus Lhx6<sup>ckO</sup> | 0.0040 | ** |
| S2E | V1 LFP power across frequency bands<br>Control versus Lhx6<sup>chHt</sup> | Control = 6<br>Lhx6<sup>chHt</sup> = 4 | Student's t-test | V1: 1-4 Hz Power<br>V1: 4-12 Hz Power<br>V1: 12-35 Hz Power<br>V1: 35-60 Hz Power<br>V1: 60-100 Hz Power | 0.4769<br>0.6483<br>0.3223<br>0.4820<br>0.6851 | ns<br>ns<br>ns<br>ns<br>ns |

<sup>a</sup>alpha = 0.05<sup>b</sup>Notation for significance levels: \*p<0.05, \*\*p<0.01, \*\*\*p<0.001, \*\*\*\* p<0.0001, ns = not significant (p>0.05)

**Table S5.** Comparison of Conditional *Mef2c* Removal on Open Field Behavior. *(Corresponding to Figure 4)*

| Figure | Comparison | N | Test<sup>a</sup> | Comparison | p-value<sup>b</sup> | Level<sup>c</sup> |
| --- | --- | --- | --- | --- | --- | --- |
| 4B | Total Tracklength | Control = 21<br>Lhx6<sup>cHet</sup> = 9<br>Lhx6<sup>cKO</sup> = 15<br>PV<sup>cKO</sup> = 7 | One-way ANOVA<br>Tukey HSD | All Groups<br>Control vs. Lhx6<sup>cHet</sup><br>Control vs. Lhx6<sup>cKO</sup><br>Control vs. PV<sup>cKO</sup><br>Lhx6<sup>cHet</sup> vs. Lhx6<sup>cKO</sup><br>PV<sup>cKO</sup> vs. Lhx6<sup>cKO</sup> | F(3,48) = 10.7,<br>p=<0.0001<br>0.6827<br><0.0001<br>0.9655<br>0.0162<br>0.0005 | ****<br>*ns*<br>****<br>*ns*<br>*<br>*** |
| 4C | Jumping | Control = 17<br>Lhx6<sup>cHet</sup> = 7<br>Lhx6<sup>cKO</sup> = 10<br>PV<sup>cKO</sup> = 7 | One-way ANOVA<br>Tukey HSD | All Groups<br>Control vs. Lhx6<sup>cHet</sup><br>Control vs. Lhx6<sup>cKO</sup><br>Control vs. PV<sup>cKO</sup><br>Lhx6<sup>cHet</sup> vs. Lhx6<sup>cKO</sup><br>Lhx6<sup>cHet</sup> vs. PV<sup>cKO</sup> | F(3,37) = 14.8,<br>p<0.0001<br>0.9936<br><0.0001<br>0.9869<br>0.0002<br><0.0001 | ****<br>*ns*<br>****<br>*ns*<br>***<br>**** |
| 4D | Tracklength by Group and Time Bin (Early, Late) | Control = 21<br>Lhx6<sup>cHet</sup> = 10<br>Lhx6<sup>cKO</sup> = 15<br>PV<sup>cKO</sup> = 6 | Two-way Repeated Measures ANOVA<br>Bonferroni | Group*Time Bin<br>Time Bin only<br>Control Early vs. Late<br>Lhx6<sup>cHet</sup> Early vs. Late<br>Lhx6<sup>cKO</sup> Early vs. Late<br>PV<sup>cKO</sup> Early vs. Late | F(3,50) = 22.5,<br>p<0.0001<br>F(1,50) = 59.4,<br>p<0.0001<br><0.0001<br><0.0001<br>0.2209<br>0.0482 | ****<br>****<br>****<br>****<br>*ns*<br>* |
| 4E | Grooming by Group and Time Bin (Early, Late) | Control = 20<br>Lhx6<sup>cHet</sup> = 10<br>Lhx6<sup>cKO</sup> = 13<br>PV<sup>cKO</sup> = 7 | Two-way Repeated Measures ANOVA<br>Bonferroni | Groups*Time Bin<br>Time Bin only<br>Control Early vs. Late<br>Lhx6<sup>cHet</sup> Early vs. Late<br>Lhx6<sup>cKO</sup> Early vs. Late<br>PV<sup>cKO</sup> Early vs. Late | F(3,46) = 6.4,<br>p=0.0010<br>F(1,46) = 16.0,<br>p=0.0002<br><0.0001<br>0.0231<br>0.9622<br>0.3653 | ***<br>***<br>****<br>*<br>*ns*<br>*ns* |

<sup>a</sup>For significant multiple comparisons (ANOVA), Tukey HSD or Bonferroni post hoc test was applied for determining pairwise differences<sup>b</sup>alpha = 0.05<sup>c</sup>Notation for significance levels: \*p<0.05, \*\*p<0.01, \*\*\*p<0.001, \*\*\*\* p<0.0001, *ns* = not significant (p>0.05)

**Table S6.** Comparison of Conditional *Mef2c* Removal on Clasping, EPM, and Y-Maze. *(Corresponding to Figure 5)*

| Figure | Comparison | N | Test<sup>a</sup> | Comparison | p-value<sup>b</sup> | Level<sup>c</sup> |
| --- | --- | --- | --- | --- | --- | --- |
| 5A | Paw Clasping | Control = 13<br>Lhx6<sup>chHet</sup> = 8<br>Lhx6<sup>ckKO</sup> = 11 | One-way ANOVA<br>Tukey HSD | All Groups<br>Control vs. Lhx6<sup>chHet</sup><br>Control vs. Lhx6<sup>ckKO</sup><br>Lhx6<sup>chHet</sup> vs. Lhx6<sup>ckKO</sup> | F (3,31) = 25.2,<br>p<0.0001<br>0.9332<br><0.0001<br><0.0001 | ****<br>*ns*<br>****<br>**** |
| 5C | EPM<br>Open Arm Time | Control = 19<br>Lhx6<sup>chHet</sup> = 16<br>Lhx6<sup>ckKO</sup> = 8 | One-way ANOVA<br>Tukey HSD | All Groups<br>Control vs. Lhx6<sup>chHet</sup><br>Control vs. Lhx6<sup>ckKO</sup><br>Lhx6<sup>chHet</sup> vs. Lhx6<sup>ckKO</sup> | F (2,40) = 10.7,<br>p=0.0002<br>0.3218<br>0.0001<br>0.0050 | ***<br>*ns*<br>***<br>** |
| 5D | EPM<br>Open Arm Entries | Control = 19<br>Lhx6<sup>chHet</sup> = 16<br>Lhx6<sup>ckKO</sup> = 8 | One-way ANOVA<br>Tukey HSD | All Groups<br>Control vs. Lhx6<sup>chHet</sup><br>Control vs. Lhx6<sup>ckKO</sup><br>Lhx6<sup>chHet</sup> vs. Lhx6<sup>ckKO</sup> | F (2,40) = 7.1,<br>p=0.0023<br>0.4494<br>0.0016<br>0.0265 | **<br>*ns*<br>**<br>* |
| 5E | EPM<br>Total Tracklength | Control = 19<br>Lhx6<sup>chHet</sup> = 16<br>Lhx6<sup>ckKO</sup> = 8 | One-way ANOVA<br>Tukey HSD | All Groups<br>Control vs. Lhx6<sup>chHet</sup><br>Control vs. Lhx6<sup>ckKO</sup><br>Lhx6<sup>chHet</sup> vs. Lhx6<sup>ckKO</sup> | F (2,40) = 5.2,<br>p=0.0083<br>0.6232<br>0.0395<br>0.0074 | *<br>*ns*<br>*<br>** |
| 5G | Y-Maze<br>Spontaneous Alternation | Control = 17<br>Lhx6<sup>chHet</sup> = 19<br>Lhx6<sup>ckKO</sup> = 8 | One-way ANOVA | All Groups | F (2,42) = 0.9,<br>p=0.4322 | *ns* |
| 5H | Y-Maze<br>Tracklength | Control = 17<br>Lhx6<sup>chHet</sup> = 19<br>Lhx6<sup>ckKO</sup> = 8 | One-way ANOVA<br>Tukey HSD | All Groups<br>Control vs. Lhx6<sup>chHet</sup><br>Control vs. Lhx6<sup>ckKO</sup><br>Lhx6<sup>chHet</sup> vs. Lhx6<sup>ckKO</sup> | F (2,42) = 14.7,<br>p<0.0001<br>0.5723<br><0.0001<br>0.0001 | ****<br>*ns*<br>****<br>*** |

<sup>a</sup>For significant multiple comparisons (ANOVA), a Tukey HSD post hoc test was applied for determining pairwise differences<sup>b</sup>alpha = 0.05<sup>c</sup>Notation for significance levels: \*p<0.05, \*\*p<0.01, \*\*\*p<0.001, \*\*\*\* p<0.0001, *ns* = not significant (p>0.05)

**Table S6 Continued. Comparison of Conditional *Mef2c* Removal on Clasping, EPM, and Y-Maze** (Corresponding to Figure 5)
| Figure | Comparison | N | Test<sup>a</sup> | Comparison | p-value<sup>b</sup> | Level<sup>c</sup> |
| --- | --- | --- | --- | --- | --- | --- |
| 5I | Y-Maze<br>Center Time | Control = 17<br>Lhx6<sup>cHet</sup> = 19<br>Lhx6<sup>cKO</sup> = 8 | One-way<br>ANOVA<br>Tukey HSD | All Groups<br>Control vs. Lhx6<sup>cHet</sup><br>Control vs. Lhx6<sup>cKO</sup><br>Lhx6<sup>cHet</sup> vs. Lhx6<sup>cKO</sup> | F (2,42) = 11.5,<br>p=0.0001<br>0.9984<br>0.0002<br>0.0002 | ***<br>*ns*<br>***<br>*** |
| 5J | Y-Maze<br>Entries | Control = 17<br>Lhx6<sup>cHet</sup> = 19<br>Lhx6<sup>cKO</sup> = 8 | One-way<br>ANOVA<br>Tukey HSD | All Groups<br>Control vs. Lhx6<sup>cHet</sup><br>Control vs. Lhx6<sup>cKO</sup><br>Lhx6<sup>cHet</sup> vs. Lhx6<sup>cKO</sup> | F (2,42) = 12.8,<br>p<0.0001<br>0.9475<br><0.0001<br>0.0001 | ****<br>*ns*<br>****<br>*** |
| 5K | Y-Maze<br>Partial Entries | Control = 17<br>Lhx6<sup>cHet</sup> = 19<br>Lhx6<sup>cKO</sup> = 8 | One-way<br>ANOVA<br>Tukey HSD | All Groups<br>Control vs. Lhx6<sup>cHet</sup><br>Control vs. Lhx6<sup>cKO</sup><br>Lhx6<sup>cHet</sup> vs. Lhx6<sup>cKO</sup> | F (2,42) = 8.98,<br>p=0.0006<br>0.7049<br>0.0035<br>0.0005 | ***<br>*ns*<br>**<br>*** |

<sup>a</sup>For significant multiple comparisons (ANOVA), a Tukey HSD post hoc test was applied for determining pairwise differences<sup>b</sup>alpha = 0.05<sup>c</sup>Notation for significance levels: \*p<0.05, \*\*p<0.01, \*\*\*p<0.001, \*\*\*\* p<0.0001, *ns* = not significant (p>0.05)

**Table S7.** Comparison of Conditional *Mef2c* Removal on Social and Cognitive Behavior. *(Corresponding to Figure 6, Figure S3)*

| Figure | Assay | N | Test<sup>a</sup> | Comparison | p-value<sup>b</sup> | Level<sup>c</sup> |
| --- | --- | --- | --- | --- | --- | --- |
| 6B | Social Preference | Control = 18<br>Lhx6<sup>cHet</sup> = 19<br>Lhx6<sup>cKO</sup> = 8 | Two-way Repeated Measures ANOVA<br>(Bonferroni) | Groups*Stimulus<br>Stimulus only<br>Control Animal vs Object<br>Lhx6<sup>cHet</sup> Animal vs Object<br>Lhx6<sup>cKO</sup> Animal vs Object | F(2,42) = 0.7,<br>p=0.5065<br>F(1,42) = 15.4,<br>p=0.0003<br>0.0181<br>0.0017<br>0.9269 | ns<br>***<br>*<br>**<br>ns |
| 6E | Novel Object Location | Control = 15<br>Lhx6<sup>cHet</sup> = 10<br>Lhx6<sup>cKO</sup> = 10<br>1 Lhx6<sup>cKO</sup> excluded due to < 3s exploration | Two-way Repeated Measures ANOVA<br>(Bonferroni) | Groups*Stimulus<br>Stimulus only<br>Control Novel vs Familiar<br>Lhx6<sup>cHet</sup> Novel vs Familiar<br>Lhx6<sup>cKO</sup> Novel vs Familiar | F(2,32) = 4.0,<br>p=0.0283<br>F(1,32) = 27.8,<br>p<0.0001<br><0.0001<br>0.0109<br>>0.9999 | *<br>****<br>****<br>*<br>ns |
| S3B | Social Interaction Bouts | Control = 7<br>Lhx6<sup>cHet</sup> = 6<br>Lhx6<sup>cKO</sup> = 5 | One-way ANOVA<br>(Tukey HSD) | All Groups<br>Control vs. Lhx6<sup>cHet</sup><br>Control vs. Lhx6<sup>cKO</sup><br>Lhx6<sup>cHet</sup> vs. Lhx6<sup>cKO</sup> | F (2,15) = 128.9,<br>p=0.0029<br>0.5870<br>0.0025<br>0.0206 | **<br>ns<br>**<br>* |
| S3C | Social Interaction Bout Duration | Control = 7<br>Lhx6<sup>cHet</sup> = 6<br>Lhx6<sup>cKO</sup> = 5 | One-way ANOVA<br>(Tukey HSD) | All Groups<br>Control vs. Lhx6<sup>cHet</sup><br>Control vs. Lhx6<sup>cKO</sup><br>Lhx6<sup>cHet</sup> vs. Lhx6<sup>cKO</sup> | F (2,15) = 6.6,<br>p=0.0089<br>0.6169<br>0.0390<br>0.0083 | **<br>ns<br>*<br>** |
| S3D | Track length in Social Interaction Assay | Control = 7<br>Lhx6<sup>cHet</sup> = 6<br>Lhx6<sup>cKO</sup> = 5 | One-way ANOVA<br>(Tukey HSD) | All Groups<br>Control vs. Lhx6<sup>cHet</sup><br>Control vs. Lhx6<sup>cKO</sup><br>Lhx6<sup>cHet</sup> vs. Lhx6<sup>cKO</sup> | F (2,15) = 19.4,<br>p<0.0001<br>0.9967<br>0.0002<br>0.0002 | ****<br>ns<br>***<br>*** |
| S3F | Novel Object Recognition | Control = 13<br>Lhx6<sup>cHet</sup> = 10<br>Lhx6<sup>cKO</sup> = 10<br>1 Lhx6<sup>cKO</sup> excluded due to < 3s exploration | Two-way Repeated Measures ANOVA | Groups*Stimulus<br>Stimulus only | F(2,30) = 0.7,<br>p=0.4973<br>F(1,30) = 2.4,<br>p=0.1325 | ns<br>ns |

<sup>a</sup>For significant multiple comparisons (ANOVA), a Bonferroni or Tukey HSD post hoc test was applied for determining pairwise differences <sup>b</sup>alpha = 0.05
<sup>c</sup>Notation for significance levels: \*p<0.05, \*\*p<0.01, \*\*\*p<0.001, \*\*\*\* p<0.0001, ns = not significant (p>0.05)

## References

1. Harrington AJ, Bridges CM, Berto S, Blankenship K, Cho JY, Assali A, et al. (2020): MEF2C Hypofunction in Neuronal and Neuroimmune Populations Produces MEF2C Haploinsufficiency Syndrome-like Behaviors in Mice. Biol Psychiatry. 88:488–499.

2. Assali A, Harrington AJ, Cowan CW (2019): Emerging roles for MEF2 in brain development and mental disorders. Curr Opin Neurobiol. 59:49–58.

3. Harrington AJ, Raissi A, Rajkovich K, Berto S, Kumar J, Molinaro G, et al. (2016): MEF2C regulates cortical inhibitory and excitatory synapses and behaviors relevant to neurodevelopmental disorders. Elife. 5.

4. Cosgrove D, Whitton L, Fahey L, P Ob, Donohoe G, Morris DW (2021): Genes influenced by MEF2C contribute to neurodevelopmental disease via gene expression changes that affect multiple types of cortical excitatory neurons. Hum Mol Genet. 30:961–970.

5. Rajkovich KE, Loerwald KW, Hale CF, Hess CT, Gibson JR, Huber KM (2017): Experience-Dependent and Differential Regulation of Local and Long-Range Excitatory Neocortical Circuits by Postsynaptic Mef2c. Neuron. 93:48–56.

6. Wan L, Liu X, Hu L, Chen H, Sun Y, Li Z, et al. (2021): Genotypes and Phenotypes of MEF2C Haploinsufficiency Syndrome: New Cases and Novel Point Mutations. Front Pediatr. 9:664449.

7. Tu S, Akhtar MW, Escorihuela RM, Amador-Arjona A, Swarup V, Parker J, et al. (2017): NitroSynapsin therapy for a mouse MEF2C haploinsufficiency model of human autism. Nat Commun. 8:1488.

8. Vormstein-Schneider D, Lin JD, Pelkey KA, Chittajallu R, Guo B, Arias-Garcia MA, et al. (2020): Viral manipulation of functionally distinct interneurons in mice, non-human primates and humans. Nat Neurosci.

9. Mayer C, Hafemeister C, Bandler RC, Machold R, Batista Brito R, Jaglin X, et al. (2018): Developmental diversification of cortical inhibitory interneurons. Nature. 555:457–462.

10. Rudy B, Fishell G, Lee S, Hjerling-Leffler J (2011): Three groups of interneurons account for nearly 100% of neocortical GABAergic neurons. Dev Neurobiol. 71:45–61.

11. Fogarty M, Grist M, Gelman D, Marín O, Pachnis V, Kessaris N (2007): Spatial genetic patterning of the embryonic neuroepithelium generates GABAergic interneuron diversity in the adult cortex. J Neurosci. 27:10935–10946.

12. Cobos I, Calcagnotto ME, Vilaythong AJ, Thwin MT, Noebels JL, Baraban SC, et al. (2005): Mice lacking Dlx1 show subtype-specific loss of interneurons, reduced inhibition and epilepsy. Nat Neurosci. 8:1059–1068.

13. Cobos I, Long JE, Thwin MT, Rubenstein JL (2006): Cellular patterns of transcription factor expression in developing cortical interneurons. Cereb Cortex. 16 Suppl 1:i82–88.

14. Alifragis P, Liapi A, Parnavelas JG (2004): Lhx6 regulates the migration of cortical interneurons from the ventral telencephalon but does not specify their GABA phenotype. J Neurosci. 24:5643–5648.

15. Lavdas AA, Grigoriou M, Pachnis V, Parnavelas JG (1999): The medial ganglionic eminence gives rise to a population of early neurons in the developing cerebral cortex. J Neurosci. 19:7881–7888.

16. Liodis P, Denaxa M, Grigoriou M, Akufo-Addo C, Yanagawa Y, Pachnis V (2007): Lhx6 activity is required for the normal migration and specification of cortical interneuron subtypes. J Neurosci. 27:3078–3089.

17. Southwell DG, Paredes MF, Galvao RP, Jones DL, Froemke RC, Sebe JY, et al. (2012): Intrinsically determined cell death of developing cortical interneurons. Nature. 491:109–113.

18. Hanson E, Armbruster M, Lau LA, Sommer ME, Klaft ZJ, Swanger SA, et al. (2019): Tonic Activation of GluN2C/GluN2D-Containing NMDA Receptors by Ambient Glutamate Facilitates Cortical Interneuron Maturation. J Neurosci. 39:3611–3626.

19. Chattopadhyaya B, Di Cristo G, Higashiyama H, Knott GW, Kuhlman SJ, Welker E, et al. (2004): Experience and activity-dependent maturation of perisomatic GABAergic innervation in primary visual cortex during a postnatal critical period. J Neurosci. 24:9598–9611.

20. del Río JA, de Lecea L, Ferrer I, Soriano E (1994): The development of parvalbumin-immunoreactivity in the neocortex of the mouse. Brain Res Dev Brain Res. 81:247–259.

21. Xu X, Roby KD, Callaway EM (2010): Immunochemical characterization of inhibitory mouse cortical neurons: three chemically distinct classes of inhibitory cells. J Comp Neurol. 518:389–404.

22. Gonchar Y, Wang Q, Burkhalter A (2007): Multiple distinct subtypes of GABAergic neurons in mouse visual cortex identified by triple immunostaining. Front Neuroanat. 1:3.

23. Lee AT, Cunniff MM, See JZ, Wilke SA, Luongo FJ, Ellwood IT, et al. (2019): VIP Interneurons Contribute to Avoidance Behavior by Regulating Information Flow across Hippocampal-Prefrontal Networks. Neuron. 102:1223-1234.e1224.

24. Allaway KC, Gabitto MI, Wapinski O, Saldi G, Wang CY, Bandler RC, et al. (2021): Genetic and epigenetic coordination of cortical interneuron development. Nature. 597:693–697.

25. Patz S, Grabert J, Gorba T, Wirth MJ, Wahle P (2004): Parvalbumin expression in visual cortical interneurons depends on neuronal activity and TrkB ligands during an Early period of postnatal development. Cereb Cortex. 14:342–351.

26. Frischknecht R, Heine M, Perrais D, Seidenbecher CI, Choquet D, Gundelfinger ED (2009): Brain extracellular matrix affects AMPA receptor lateral mobility and short-term synaptic plasticity. Nat Neurosci. 12:897–904.

27. Sigal YM, Bae H, Bogart LJ, Hensch TK, Zhuang X (2019): Structural maturation of cortical perineuronal nets and their perforating synapses revealed by superresolution imaging. Proc Natl Acad Sci U S A. 116:7071–7076.

28. Fawcett JW, Fyhn M, Jendelova P, Kwok JCF, Ruzicka J, Sorg BA (2022): The extracellular matrix and perineuronal nets in memory. Mol Psychiatry. 27:3192–3203.

29. Ruzicka J, Dalecka M, Safrankova K, Peretti D, Jendelova P, Kwok JCF, et al. (2022): Perineuronal nets affect memory and learning after synapse withdrawal. Transl Psychiatry. 12:480.

30. Barbosa AC, Kim MS, Ertunc M, Adachi M, Nelson ED, McAnally J, et al. (2008): MEF2C, a transcription factor that facilitates learning and memory by negative regulation of synapse numbers and function. Proc Natl Acad Sci U S A. 105:9391–9396.

31. Flavell SW, Cowan CW, Kim TK, Greer PL, Lin Y, Paradis S, et al. (2006): Activity-dependent regulation of MEF2 transcription factors suppresses excitatory synapse number. Science. 311:1008–1012.

32. Mitchell AC, Javidfar B, Pothula V, Ibi D, Shen EY, Peter CJ, et al. (2018): MEF2C transcription factor is associated with the genetic and epigenetic risk architecture of schizophrenia and improves cognition in mice. Mol Psychiatry. 23:123–132.

33. Vinck M, Batista-Brito R, Knoblich U, Cardin JA (2015): Arousal and locomotion make distinct contributions to cortical activity patterns and visual encoding. Neuron. 86:740–754.

34. Gao R, Peterson EJ, Voytek B (2017): Inferring synaptic excitation/inhibition balance from field potentials. NeuroImage. 158:70–78.

35. Donoghue T, Schaworonkow N, Voytek B (2022): Methodological considerations for studying neural oscillations. Eur J Neurosci. 55:3502–3527.

36. Li H, Radford JC, Ragusa MJ, Shea KL, McKercher SR, Zaremba JD, et al. (2008): Transcription factor MEF2C influences neural stem/progenitor cell differentiation and maturation in vivo. Proc Natl Acad Sci U S A. 105:9397–9402.

37. Kong Y, Zhou W, Sun Z (2020): Nuclear receptor corepressors in intellectual disability and autism. Mol Psychiatry. 25:2220–2236.

38. Adachi M, Lin PY, Pranav H, Monteggia LM (2016): Postnatal Loss of Mef2c Results in Dissociation of Effects on Synapse Number and Learning and Memory. Biol Psychiatry. 80:140–148.

39. Schmeisser MJ, Ey E, Wegener S, Bockmann J, Stempel AV, Kuebler A, et al. (2012): Autistic-like behaviours and hyperactivity in mice lacking ProSAP1/Shank2. Nature. 486:256–260.

40. Norwood J, Franklin JM, Sharma D, D’Mello SR (2014): Histone deacetylase 3 is necessary for proper brain development. J Biol Chem. 289:34569–34582.

41. El Hayek L, Tuncay IO, Nijem N, Russell J, Ludwig S, Kaur K, et al. (2020): KDM5A mutations identified in autism spectrum disorder using forward genetics. Elife. 9.

42. Nguyen MV, D. F, Felice CA, Shan X, Nigam A, Mandel G, et al. (2012): MeCP2 is critical for maintaining mature neuronal networks and global brain anatomy during late stages of postnatal brain development and in the mature adult brain. J Neurosci. 32:10021–10034.

43. Kriaucionis S, Bird A (2003): DNA methylation and Rett syndrome. Hum Mol Genet. 12 Spec No 2:R221–227.

44. Moy SS, Nadler JJ, Young NB, Perez A, Holloway LP, Barbaro RP, et al. (2007): Mouse behavioral tasks relevant to autism: phenotypes of 10 inbred strains. Behav Brain Res. 176:4–20.

45. Crawley JN (2004): Designing mouse behavioral tasks relevant to autistic-like behaviors. Ment Retard Dev Disabil Res Rev. 10:248–258.

46. Crespi B (2021): Pattern Unifies Autism. Front Psychiatry. 12:621659.

47. Paterno R, Marafiga JR, Ramsay H, Li T, Salvati KA, Baraban SC (2021): Hippocampal gamma and sharp-wave ripple oscillations are altered in a Cntnap2 mouse model of autism spectrum disorder. Cell Rep. 37:109970.

48. Pagano J, Landi S, Stefanoni A, Nardi G, Albanesi M, Bauer HF, et al. (2023): Shank3 deletion in PV neurons is associated with abnormal behaviors and neuronal functions that are rescued by increasing GABAergic signaling. Mol Autism. 14:28.

49. Cooley Coleman JA, Sarasua SM, Boccuto L, Moore HW, Skinner SA, DeLuca JM (2021): Comprehensive investigation of the phenotype of MEF2C-related disorders in human patients: A systematic review. Am J Med Genet A. 185:3884–3894.

50. Cooley Coleman JA, Sarasua SM, Moore HW, Boccuto L, Cowan CW, Skinner SA, et al. (2022): Clinical findings from the landmark MEF2C-related disorders natural history study. Mol Genet Genomic Med. 10:e1919.

51. Chaudhary R, Agarwal V, Kaushik AS, Rehman M (2021): Involvement of myocyte enhancer factor 2c in the pathogenesis of autism spectrum disorder. Heliyon. 7:e06854.

52. Fahey L, Ali D, Donohoe G, P ÓB, Morris DW (2023): Genes positively regulated by Mef2c in cortical neurons are enriched for common genetic variation associated with IQ and educational attainment. Hum Mol Genet. 32:3194–3203.

53. Pai EL, Chen J, Fazel Darbandi S, Cho FS, Chen J, Lindtner S, et al. (2020): Maf and Mafb control mouse pallial interneuron fate and maturation through neuropsychiatric disease gene regulation. Elife. 9.

54. Sohal VS, Rubenstein JLR (2019): Excitation-inhibition balance as a framework for investigating mechanisms in neuropsychiatric disorders. Mol Psychiatry. 24:1248–1257.

55. Li W-K, Zhang S-Q, Peng W-L, Shi Y-H, Yuan B, Yuan Y-T, et al. (2023): Whole-brain in vivo base editing reverses behavioral changes in Mef2c-mutant mice. Nature Neuroscience.

## Supplemental References

1. Paxinos GaF K.B.J. (2001): The Mouse Brain in Stereotaxic Coordinates. 2nd ed. San Diego: Academic Press

2. Vormstein-Schneider D, Lin JD, Pelkey KA, Chittajallu R, Guo B, Arias-Garcia MA et al. (2020): Viral manipulation of functionally distinct interneurons in mice, non-human primates and humans. Nat Neurosci.

